# The essential lipoprotein maturation enzyme Lgt is required for flagellar and membrane integrity in *Helicobacter pylori*

**DOI:** 10.64898/2026.10.09.757914

**Authors:** Tristan Ruffiot, Samia Hicham, Ana Oliveira Paiva, Bertrand Legeret, Dynang Seng, Yonghua Li-Beisson, Ivo Gomperts Boneca, Nienke Buddelmeijer

**Affiliations:** Institut Pasteur, Université Paris Cité, CNRS UMR6047, INSERM U1306, Unité Biologie et Génétique de la Paroi Bactérienne, Paris, FRANCE; HelioBiotec Marseille CEA, BIAM UMR 7265, FRANCE

## Abstract

Pre-prolipoprotein phosphatidylglycerol diacylglyceryl transferase (Lgt) catalyzes the first step in the lipoprotein modification pathway and is essential for growth and viability of *Helicobacter pylori*. We explored the fate of *H. pylori* cells under Lgt depletion conditions to obtain insight into Lgt function on morphology and physiology. Downregulation of *lgt* expression led to growth arrest and a heterogenous population of small rod-shaped cells and small round cells with spiky protrusions, a loose outer membrane and disintegrated flagella. *H. pylori* overcomes a defect in lipoprotein modification through alteration of its gene expression profile resulting in so-called revertants. Outer membrane protein synthesis, carbon metabolism and nutrient uptake was reduced, and protein translation and flagellar biogenesis increased. Lgt specific regulation was observed in phospholipid biosynthesis resulting in an initial increase in cardiolipin followed by an increase in lysophospholipids at the time of adaptation. Suppressors of Δ*lgt* that were selected on solid media under restrictive conditions either lacked chromosomal mutations or had mutations in the *cag*UT-*tet*R regulon controlling the expression of *lgt*. A suppressor without mutations had an altered expression profile that led to downregulation of a DNA methyltransferase and upregulation of HP_0014, a motility associated glycosyltransferase family protein. Transmission electron microscopy and single cell video microscopy showed that a subpopulation of cells is motile in the revertant while the suppressor regained full motility with higher average speed compared to the wild type strain. Overall, our data show that lipoprotein modification by Lgt is important for membrane and flagellar integrity and that the heterogenous character of *H. pylori* cultures allows individual cells to overcome this cell envelope stress and survive. Our findings are important for antibacterial development strategies targeting essential integral membrane enzymes involved in lipoprotein modification and stimulate exploration of synergetic activity between known antibiotics and novel antibacterial agents targeting novel cellular pathways.

**Importance:** Studying essential proteins on the fate of their absence is challenging since it requires controlled expression systems that can be regulated. Low levels of Lgt greatly impact membrane and flagellar integrity disturbing cell envelope function. *H. pylori* is well known for its genomic plasticity and very efficiently circumvents various threats. This work describes how *H. pylori* overcomes deleterious effects caused by incomplete maturation of lipoproteins and suggests that inhibition of Lgt leads to a bacteriostatic effect in *H. pylori*. These results should be considered in the development of novel antibacterial agents targeting Lgt and related enzymes.

## Introduction

*Helicobacter pylori* is a pathogen that colonizes the human stomach causing gastritis and peptic ulcers that, when left untreated, may lead to stomach cancer. As seen with many bacterial species, the rise in antibiotic resistance is also a concern with *H. pylori* [1], even though this bacterium is no longer listed as a priority pathogen by the WHO [2]. *H. pylori* has a helical rod cell shape that, together with its polar flagella, allows for high-speed motility and rapid penetration of the mucus layer [3]. It also secretes urease that converts urea into carbon dioxide and ammonium, resulting in a local increase in pH, reducing the density of the mucus layer [4]. Subsequently, *H. pylori* adheres to epithelial cells via its flagella and adhesins and secretes various virulence factors into host cells to successfully colonize its niche [5]. Under stress conditions, such as elevated oxygen levels or nutrient deprivation, *H. pylori* changes shape from a rod-helical into a spherical coccoid cell [6,7]. Coccoid cells are viable, undergo limited DNA replication [8–10] and maintain a basic level of metabolic activity [8,11], however, a process to revert coccoid cells back to helical-rod shape is still a topic of debate [12].

Bacterial lipoproteins fulfil important functions in the cell envelope as part of essential membrane protein machineries and are potent virulence factors. Lipoproteins of diderm (two membrane) bacteria are modified at their amino termini by fatty acids through the lipoprotein modification pathway composed of three integral membrane enzymes [13]. The first and committed step is catalyzed by pre-prolipoprotein phosphatidylglycerol diacylglyceryl transferase (Lgt) that adds a diacylglyceryl moiety from phosphatidylglycerol, a membrane phospholipid, onto the conserved cysteine in the recognition site (lipobox) of pre-prolipoproteins. Signal peptidase II (Lsp) then cleaves the signal peptide, resulting in prolipoprotein where diacylglyceryl-cysteine becomes the first amino acids in the protein sequence. Finally, lipoprotein N-acyltransferase (Lnt) adds a third fatty acid onto the amine group of diacylglyceryl-cysteine, resulting in mature triacylated lipoprotein. Lgt and Lsp but not Lnt are essential for growth and viability of *H. pylori* [14], however, Lnt is essential for stomach colonization [15]. We recently reported that Lgt is structurally and functionally conserved in pathogenic bacteria, and that the least conserved (exposed) domains of the protein determine species specificity [16]. Lgt of *H. pylori* complements an Lgt depletion strain of *E. coli* [14,16], while it only shares 30% identity with Lgt of *E. coli* and has a smaller periplasmic domain [16]. Determination of Lgt structure [17], computational modeling of its reaction mechanism [18] and analysis of conserved residues and functional domains [16,17,19] led to the hypothesis that Lgt may be a promising target for the development of novel antibiotics. Indeed, Lgt was identified as target of a small inhibitory molecule in *E. coli* by a phenotypic screen for inhibitors [20] and shown to be inhibited in an *in vitro* assay by a synthetic cyclic peptide [21].

In diderm bacteria, lipoproteins are translocated to the outer membrane by the lipoprotein outer membrane localization (Lol) machinery composed of an ABC transporter (LolCDE), a periplasmic chaperone (LolA) and an OM receptor (LolB). Some bacteria, including *H. pylori*have a slightly different ABC transporter; a LolF dimer instead of LolEC that functions together with ATPase LolD in the release of lipoproteins from the cytoplasmic membrane [22]. *H. pylori* lacks a LolB homolog [14]. Lipoproteins stimulate the innate immune system through interaction of their lipid moiety with Toll-like receptors (TLR2) where diacylated lipoproteins are recognized by TLR2/6 and triacylated lipoproteins by TLR2/1 [23]. Lipoproteins in *H. pylori* play an important role in adhesion [24], virulence [25], competence [26] and host cell migration [27]. More recent findings reported the implication of lipoproteins in cell envelope stability and flagellar function [28,29]. FapH (HP0838) is a flagellar accessory protein with a role in outer membrane ring formation conserving the OM barrier function during flagellar rotation [30]. The effect on motility and antibiotic sensitivity in mutants defective in individual lipoproteins, suggests that this class of proteins is essential for *H. pylori* pathogenicity.

Here we investigated the role of lipoprotein modification enzyme Lgt on growth, morphology and viability of *H. pylori* with the goal to use it as target for the development of novel antibiotics. Our results show that *H. pylori* alters its gene expression profile to restore flagellar function and membrane integrity to overcome a cell envelope stress caused by depletion of Lgt but without complete restoration of a wild type phenotype. The findings strengthen the idea that Lgt is a promising antibacterial target in *H. pylori*.

## Results

### Depletion of Lgt leads to growth arrest, a decrease in viability and altered cell morphology in *H. pylori*

To investigate the role of lipoprotein modification on physiology and pathogenicity of *H. pylori*, we constructed a Lgt depletion strain in N6 background similar to the procedure described by McClain *et al*. [14]. In this strain the only copy of *lgt* is under the control of a TetR regulated CagUT promoter (P*_cagUT_*) positioned between genes HP_0630 and HP_0631 (Fig. 1A). *TetR* is integrated into the *ureA* gene and constitutively expressed from the *ureA* promoter. The addition of anhydrotetracycline (aTc) results in release of TetR from P*_cagUT_* and consequent expression of *lgt*. Endogenous *lgt* (HP_0955) was replaced by insertion of a gentamycin resistance cassette. For clarity we use Δ*lgt* + and Δ*lgt* - nomenclature for Lgt depletion strain TR03 (Table S1) throughout the text to indicate expression of *lgt* in the presence of aTc and Lgt depletion in the absence of inducer, respectively. Growth of Δ*lgt* + is similar to wild type (Fig. 1B). Depletion of Lgt results in growth arrest up to approximately 15 generations (60h) when re-growth is observed (so-called revertant) (Fig. 1B). A decrease in cell viability (CFU/mL) is observed at 60h in wild type and Δ*lgt* + (Fig. 1C) due to the formation of coccoid cells (Fig. 2). Depletion of Lgt leads to a significant reduction in viability, although not below 10^3^ CFU/mL, which is re-established at the time of recurrent growth (Fig. 1C, Fig. S1A). These results suggest that *H. pylori* is able to overcome deleterious effects of Lgt depletion on a short timescale when grown in liquid medium.

**Fig. 1.**
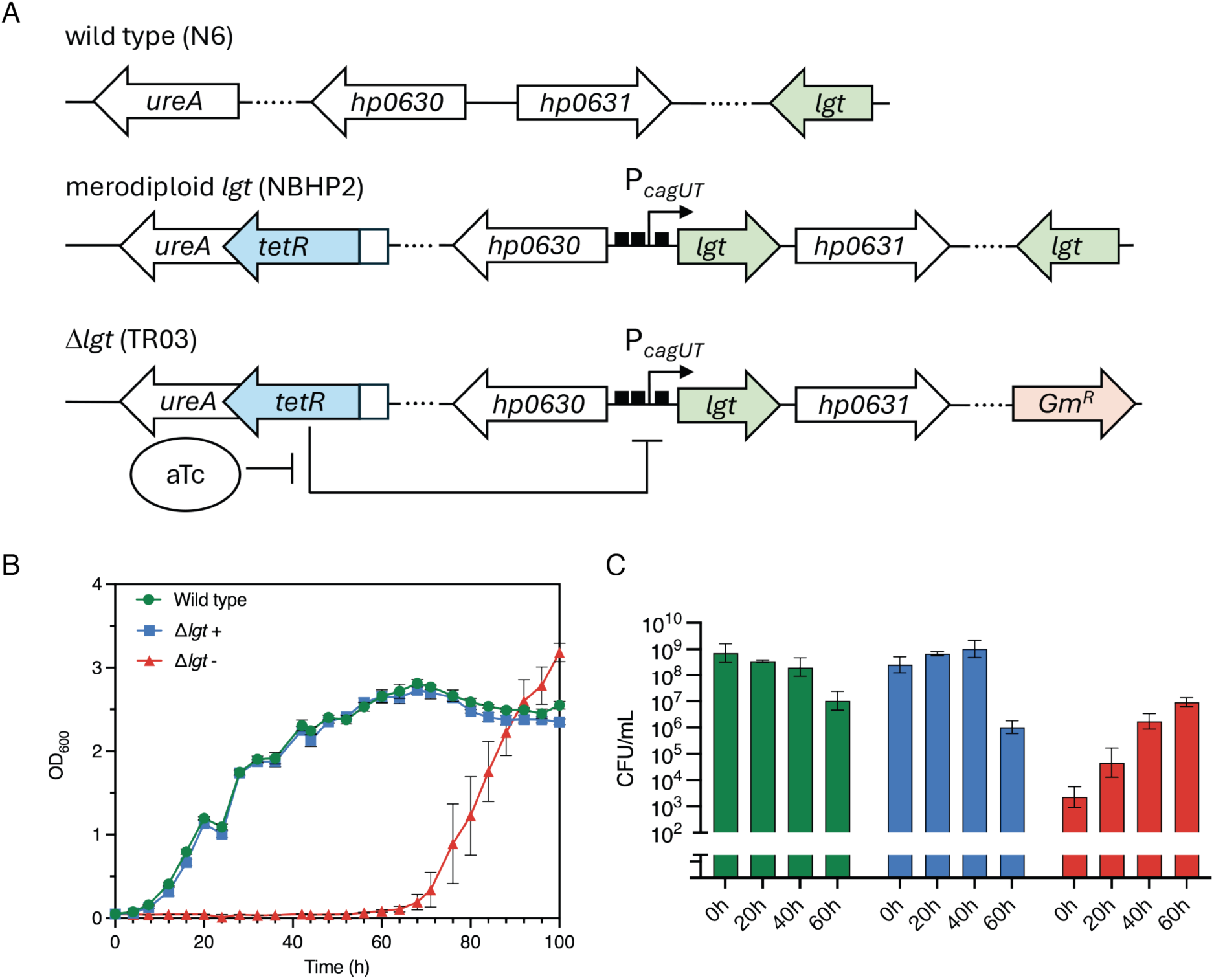
Appearance of Δ*lgt* revertants upon extended depletion of Lgt in *H. pylori*. A) Construction of a Lgt depletion strain in wild type *H. pylori* N6 by creation of an intermediate merodiploid strain with a second copy of *lgt* under control of a TetR regulated *cagUT* promoter at a neutral locus between genes *hp0630* and *hp0631* as described by McClain *et al.* [14]. Endogenous *lgt* (HP_0955) was replaced by a *Gm^R^* resistance cassette in Δ*lgt* that is now dependent on inducer (aTc) for growth. B) Growth as measured by OD_600_ of wild type and Δ*lgt* with (+) and without (-) aTc in 100 mL BHI at 37°C under microaerophilic conditions cultured in flasks. C) Viability measured as CFU/mL at time points corresponding to growth curves shown in panel B. Experiments were performed in triplicate. Spot dilution assay is shown in Fig. S1A.

**Fig. 2.**
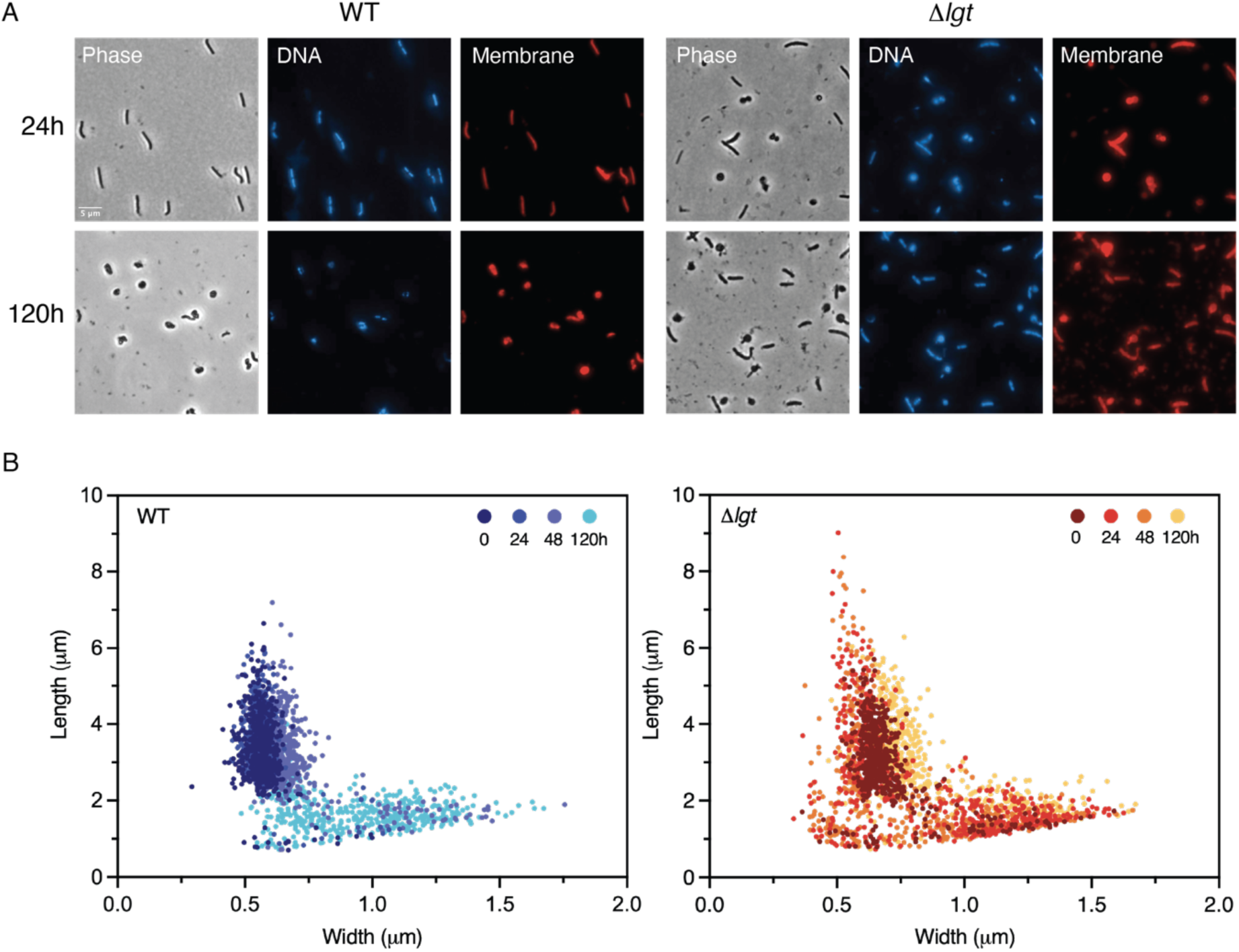
Depletion of Lgt results in a heterogenous cell population with an aberrant cell morphology in *H. pylori*. A) Epifluorescence microscopy analysis of fixed cells from a wild type and Δ*lgt* strain of *H. pylori* grown in BHI and harvested at exponential (24h) and late stationary phase (120h) (corresponding growth curves are shown in Fig. S1B). Strain Δ*lgt* was grown in the absence of inducer aTc to deplete Lgt. Nucleoids (DNA) were stained with Hoechst, and membranes were stained with FM4-64X. Scale bar is 5 μm. B) Quantification of cell parameters were performed using the MicrobeJ plugin in Fiji from two biological replicates with 500-1000 cells per condition. Evolution of cell shape at intermediate time points is presented in Fig. S1C.

At exponential phase (24h), cells of a wild type strain are rod shape (Fig. 2A) with most cells possessing two nucleoids demonstrating active DNA replication (Fig. S1A and B). At late stationary phase (120h) most cells are coccoid with only a few Hoechst-stained cells visualizing DNA (Fig. 2A). Under complete Lgt depletion conditions, Δ*lgt* forms short rods and round cells at 24h and a mixed population of rod and coccoid at 120h (Fig. 2A). To obtain more details on cell morphology changes in the population, cell parameters were measured on phase contrast images, and cell length was plotted against cell width (Fig. 2B). In the wild type strain distinct clusters can be distinguished. Cells from a 24h culture are long, slender and rod-shaped; at 48h cells start to transition from rod to coccoid; and cells from a 120h culture are as long as wide, exhibiting the characteristics of a sphere corresponding to their coccoid cell shape. Over time, cells increase in width but decrease in length before becoming shorter and wider (Fig. S1C). Depletion of Lgt results in a largely heterogenous cell population, which seems to be most diverse after 24h of growth; cells are both long and thin and short and wide (Fig. S1C), with rod-shaped cells possessing up to four nucleoids (Fig. S1C). At 120h, the population resembles wild type at 48h, suggesting that the ultimate morphology is similar between the two strains. Altogether, the results show that Lgt is essential for growth, viability and maintenance of rod cell shape in *H. pylori*.

### Adaptation of transcription profile to cell envelope stress caused by Lgt depletion

The observation that *H. pylori* can adapt to prolonged depletion of Lgt through the appearance of revertants is intriguing. We described a similar phenomenon in *E. coli*, where depletion of Lgt also led to the appearance of revertants [31]. Compensatory chromosomal mutations were not identified in *E. coli* but the revertants depended on the presence of the complementation plasmid carrying the *lgt* gene. These findings suggested that Lgt depletion did not result in a complete loss of enzyme and that low levels of *lgt* expression together with additional adaptations resulted in restored growth [31].

We performed whole genome sequencing (WGS) on the Δ*lgt* revertant of *H. pylori* in parallel with wild type N6, the merodiploid *lgt* strain and Δ*lgt* +. All genetic modifications to create the Lgt depletion strain were confirmed, including TetR operator sites surrounding P*_cagUT_*-*lgt* but no mutations were observed in the Δ*lgt* revertant that could explain its capacity for regained growth. We next selected suppressors of Δ*lgt* by growing strain TR03 on solid medium in the absence of inducer for 4 days. WGS of three selected colonies showed that two clones carried mutations in the TetR gene resulting in a premature stop codon (Table S2), likely reducing TetR binding leading to an increase in *lgt* expression. The third clone did not contain mutations (Table S2).

We therefore performed transcriptome analyses to elucidate the underlying mechanism of adaptation to Lgt depletion in *H. pylori*. Attempts to isolate RNA from Δ*lgt* - were unsuccessful, likely due to disintegration of cells with loss of nucleotides. RNASeq analyses were performed on wild type, merodiploid *lgt*, Δ*lgt* +, the Δ*lgt* revertant, and the Δ*lgt* suppressor. The *lgt* expression level is higher in the merodiploid strain and the Δ*lgt* suppressor and lower in Δ*lgt* + compared to wild type (Table S3). In the Δ*lgt* revertant, *lgt* transcript levels are much lower compared to the wild type, the merodiploid strain and the Δ*lgt* suppressor but higher compared to Δ*lgt* + (Table S3). Finally, *lgt* expression in the Δ*lgt* suppressor is higher compared to all other strains. This shows that restoration of growth and viability upon Lgt depletion leads to at least an increase in expression level of *lgt*. We further compared *tet*R expression levels in the different strains and conditions to exclude a general defect in *lgt* expression different from specific changes in the revertant causing an altered phenotype. The level of *tet*R expression is similar in the merodiploid, Δ*lgt* + and revertant, indicating that expression of *tet*R from the *ure*A promoter is not affected, however, the level is slightly higher in the suppressor (Table S4).

Comparison of the merodiploid strain with the wild type strain and the merodiploid strain with Δ*lgt* + show that the expression of several genes is affected, while wild type and Δ*lgt* + are similar (Table S4). Genes shown down- or upregulated in the merodiploid strain were inversely regulated in Δ*lgt* + (Table S4).

Using a cut-off of 3-fold change in differential gene expression, a total of 66 genes were downregulated, and 48 genes upregulated in the revertant compared to Δ*lgt* + (Fig. 3A), which we manually annotated for cellular localization (Fig. 3B) and function (Fig. 3C). Globally, outer membrane proteins, transport associated proteins and ATP associated proteins were downregulated, and ribosomal proteins, flagella associated proteins and methyl associated proteins were upregulated. Genes encoding proteins directly involved in phospholipid biosynthesis (Fig. S2) were either downregulated; *cdh* (HP_0871), or upregulated; *pgsA* (HP_1016) that, as a consequence, would increase the level of phosphatidylglycerol, the substrate of Lgt.

**Fig. 3.**
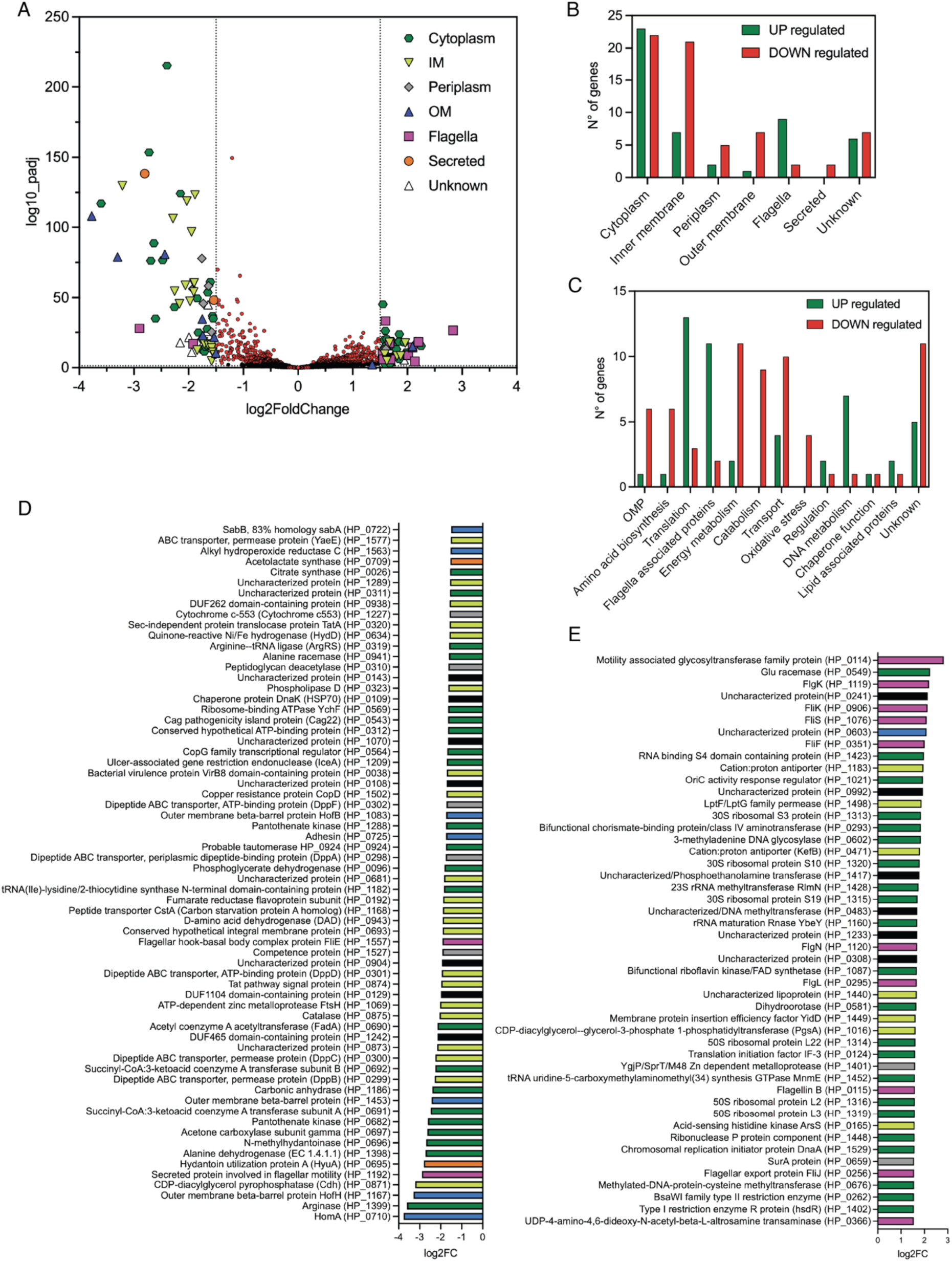
Specific transcriptome profile in Δ*lgt* revertant of *H. pylori*. A) Volcano plot representing differential gene expression as determined by RNA seq of Δ*lgt* revertant compared to Δ*lgt* induced with aTc. The vertical dotted lines represent the 1.5x log2Fold Change threshold; genes with values lower than 1.5x are indicated in red. The false discovery rate (FDR) corresponds to a p-value < 0.05 (not shown). Colors correspond to the assigned cellular localization of proteins: cytoplasm, inner membrane (IM), periplasm, outer membrane (OM), flagellar component or regulator (flagella), secreted and unknown. B) Cellular localization of gene products differentially expressed corresponding to panel A. C) Number of genes differentially expressed and classed according to functional annotation. D) List of downregulated genes and encoded proteins in the Δ*lgt* revertant relative to Δ*lgt* +. E) List of upregulated genes and encoded proteins in the Δ*lgt* revertant relative to Δ*lgt* +. Colors correspond to cellular localization depicted in panel A. Gene numbers refer to the genome of *H. pylori* strain 26695. Experiments were performed with 4 biological replicates.

Among the downregulated genes we identified three major pathways: ABC transporters, secretion systems and catabolism (Fig. 3D). The transporters include dipeptide ABC transport complex (*dppA* (HP_0298), *dppB* (HP_0299), *dppC* (HP_0300), *dppD* (HP_0301 and *dppF* (HP_0302)) and carbon transporter component *cstA* (HP_1168). Both Tat and Type IV secretion components were identified. Catabolism pathways include coenzyme A synthesis and global energy metabolism (*fadA* (HP_0690), *scoA* (HP_0691), *scoB* (HP_0692) *frdA* (HP_0192), pantothenate kinase (HP_0682), *hydD* (HP_1288), cytochrome c-553 (HP_1227), and *gltA* (HP_0026)). Six genes encoding OMPs and adhesins were also found to be downregulated (*homA* (HP_0710), *hofH* (HP_1167), HP_1453, *sabA* (HP_0725), *hofB* (HP_1083) and *sabB* (HP_0722). Critical flagellar gene *fliE* (HP_1557), a component of the MS ring-rod linker [32], was downregulated, where several other genes related to flagellar biogenesis are upregulated (Fig. 3D, Table S4).

Among the most significantly upregulated genes are those encoding ribosome subunits and tRNAs, ribonuclease gene *rnpA* (HP_1448), and translation initiation factor *infC* (HP_0124) (Fig. 3E). Together a total of 17 (31%) upregulated genes is implicated in protein translation. We also identified 11 (20%) genes involved in motility (*pseC* (HP_0366), *fliJ* (HP_0256), *flaB* (HP_0115), and HP_0295 (encoding a FlaB homolog), *flgN* (HP_1120), *fliF* (HP_), *fliS* (HP_0753), *fliK* (HP_0906), *flgK* (HP_1119), and HP_0114 that were upregulated. Finally, 6 upregulated genes are involved in DNA metabolism and DNA methylation (HP_1402, *hpyA* (HP_0262), HP_0676, *dnaA* (HP_1529), HP_0483 and HP_0602) (Fig. 3E). Overall, 58% of downregulated genes and 42% of upregulated genes encode membrane and surface proteins and more than 10 downregulated genes and 5 upregulated genes are still uncharacterized. The results indicate that revertants express *lgt* at low level that together with an altered transcription profile of important cell envelope components are able to grow.

The transcription profile of the Δ*lgt* suppressor lacking mutations in P*_cagUT_*/TetR operator sites shows a higher expression level of *lgt* (Table S3), 15 genes were downregulated and 8 genes were upregulated compared to wild type *H. pylori* (Table S4). The downregulated genes encode outer membrane proteins (*hofB* (HP_1083), *hofH* (HP_1167), *horK* (HP_1501), *babB* (HP_0896) and *frpB* (HP_1512)), proteins involved in Type IV secretion (*virB4*/*comB4* (HP_0017), *virB10/comB10* (HP_0042)), and in DNA methylation (type III methyltransferase (HP_1369)). *HofB* and *hofH* were also downregulated in the revertant. The upregulated genes are *fecA* (HP_0907), HP_0057 (Lpp20-like), a predicted lipoprotein, *pabB* (HP_0293), HP_0114 and 4 uncharacterized genes. HP_0114 is also upregulated in the Δ*lgt* revertant.

Short-term changes are observed in the Δ*lgt* revertant, while long-term changes are observed in the Δ*lgt* suppressor. Overall, *H. pylori* compensates for deleterious effects caused by Lgt depletion by altering its membrane composition, reducing transport and catabolic processes and increasing protein synthesis and flagellar biogenesis.

### Lgt depletion alters the phospholipid composition of membranes

Lgt depletion affects the expression of at least two important genes in phospholipid biosynthesis, suggesting an alteration in membrane composition under these conditions. We determined the phospholipid composition of membranes by Ultra-Performance Liquid Chromatography-Tandem Mass Spectrometry (UPLC-MS/MS) and gas-chromatography coupled to flame ionization detection and mass spectrometry (GC-FID-MS) [33,34]. The main phospholipid species in *H. pylori* membranes are phosphatidylethanolamine (PE), phosphatidylglycerol (PG), cardiolipin (CL) and phosphatidylcholine (PC) with myristate (C14:0) as main fatty acid species (Table S5).

The total amount of each phospholipid class in each strain and condition was compared to the wild type strain (Fig. 4A). Δ*lgt* + contained slightly more PG, LPG and LPC and less LPE, while depletion of Lgt (Δ*lgt* -) led to a slight increase in PE and a substantial increase in CL, PC and LPC. In both the Δ*lgt* revertant and suppressor, levels of PG, LPG and LPC are higher while PE, CL and PC levels are lower compared to wild type. In the Δ*lgt* revertant LPE levels are also higher. Based on our transcriptome data, the conversion of cytidine diphosphate-diacylglycerol (CDP-DAG) into phosphatidylglycerophosphate (PGP) by PgsA (Fig. S2) is predicted to be higher, and through downregulation of *cdh*, the level of diacylglycerol-3- phosphate lower, resulting in more PG in the Δ*lgt* revertant, which was indeed observed. In the Δ*lgt* suppressor *cdh* is also downregulated but levels of PG and LPG are slightly lower than in the Δ*lgt* revertant suggesting additional modification of PG in this mutant. Strikingly, the level of lysophospholipids, in particular LPG and LPC, is elevated in both the Δ*lgt* revertant and suppressor, suggesting increased phospholipase activity in these mutants resulting in an overall increase in membrane fluidity.

**Fig. 4.**
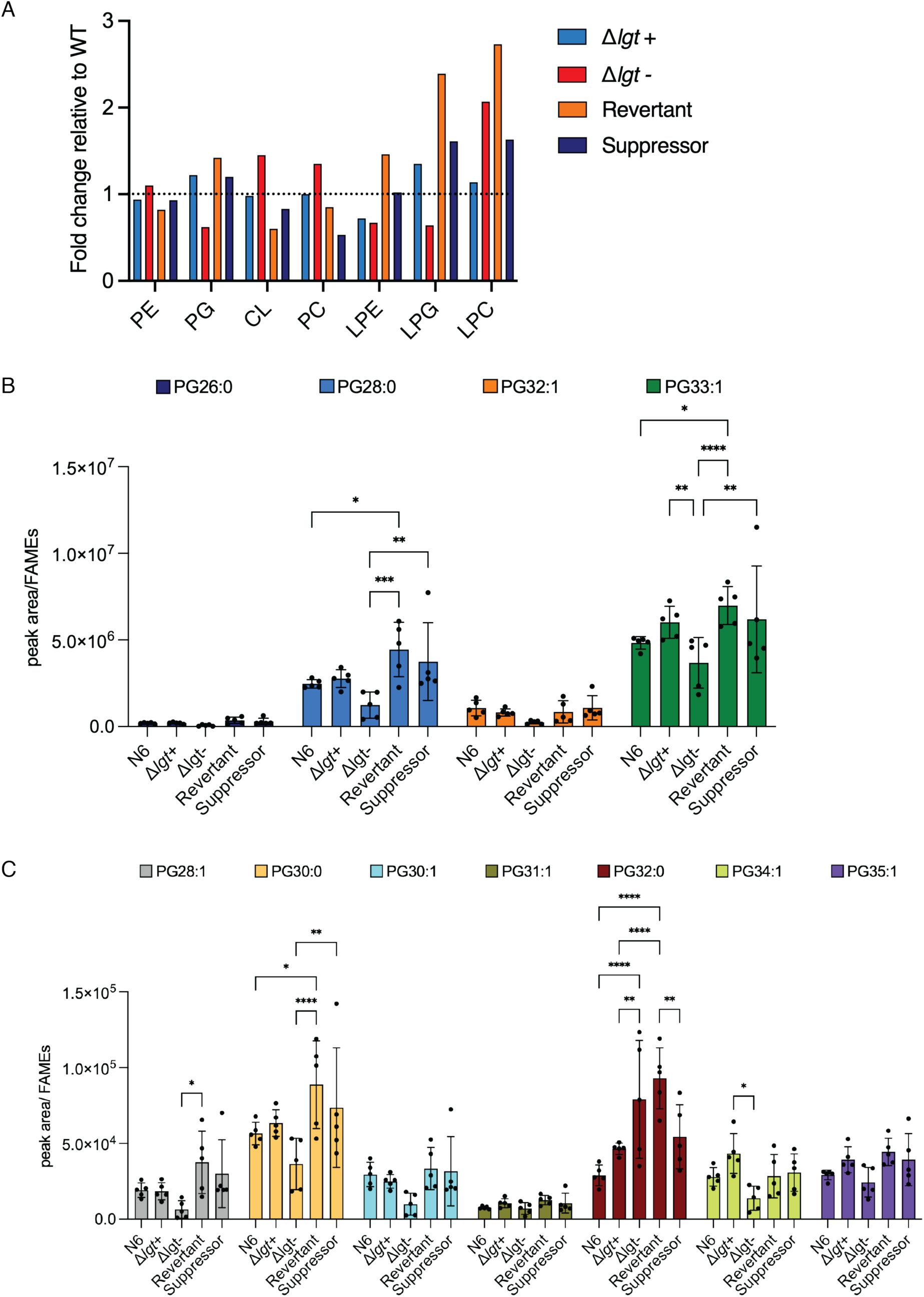
Lgt depletion alters phospholipid composition of *H. pylori* membranes. A) Quantification of phospholipid classes in membranes of Δ*lgt*, revertant and suppressor relative to wild type *H. pylori* N6. B) Quantity of major PG species relative to total amount of fatty acids. C) Quantity of minor PG species relative to total amount of fatty acids. The amount of phospholipid was calculated from the peak area from UPLC-MS analyses relative to the FID trace of FAMEs. Analysis was performed with data from five biological replicates. Statistical analysis was performed by two-way ANOVA test with * p-value <0.05, ** p-value <0.01, *** p-value <0.001 and **** p-value <0.0001. Non-significant changes (p-value > 0.05) are not shown. Data were processed using Prism 10.4.1 (GraphPad).

McClain *et al*. [35] showed that model lipoproteins CagT and Lpp20 are modified by palmitate (C16:0) and stearic acid (C18:0). We therefore characterized the fatty acid moiety of each phospholipid class to determine a correlation between phospholipid class and species in lipoprotein modification (Table S5). The main phospholipid species in PG, the acyl donor in the Lgt reaction, are PG26:0, PG28:0, PG32:1 and PG33:1 (Fig. 4B). The minor species are PG28:1, PG30:0, PG30:1, PG31:1, PG32:0, PG34:1 and PG35:1 (Fig. 4C). The amount of PG33:1 is decreased under Lgt depletion conditions (Fig. 4B). Among the minor PG species an increase is observed in PG32:0 and a decrease is observed in PG34:1(Fig. 4C). In both the revertant and suppressor main species PG28:0 and PG33:1 and minor species PG30:0 are increased relative to the depletion condition (Fig. 4B and C). Finally, the amount of PG32:0 is lower in the suppressor compared to the revertant (Fig. 4C).

The main PE species in *H. pylori* are similar to PG (Fig. S3A). Contrary to PG levels, the amount of PE33:1 is slightly higher in the Lgt-depleted condition (Fig. S3A). Among the minor PE species an increase in PE32:0, as well as a slight increase in PE30:0, is observed under Lgt depletion conditions, as seen for PG32:0 (Fig S3B). It is important to note that phospholipid quantities vary between the biological replicates in particular in Δ*lgt* -, reflecting the heterogenic character of the mutant.

The main CL species is CL61:1, which is increased in Δ*lgt* - and lower in the revertant and suppressor similar to wild type (Fig. S4A). In PC, PC32:1 and PC34:1 are the main species, for which significant changes are observed in the revertant and suppressor (Fig. S4B). Finally, the increase in lysophospholipid, particularly in the revertant, is caused by PE14:0 and PE19:0 but not PE18:1 (Fig. S4C). Furthermore, no species-specific changes are observed for LPC, and LPG is only present as LPG19:0, which is overall present in low quantities in *H. pylori* (Fig. S4C).

Changes in total amount of fatty acids were also observed. An increase in C16:0, C18:0 and C18 cyclo is observed in Δ*lgt* - that are again lower in the Δ*lgt* revertant and suppressor mutants (Fig. S5A). Differences in minor fatty acids; C12:0, C14:1, C16:1, C18cyclo2, C18cyclo3, C18cyclo4 and C18cyclo5, were not significant (Fig. S5B). Together the results show that the phospholipid composition drastically changes when lipoprotein modification is disturbed.

### Disruption of flagellar integrity caused by Lgt depletion leads to a defect in motility

The variable expression level of a large number of genes involved in flagella biogenesis suggests that Lgt depletion impacts flagellar function and motility. Furthermore, cardiolipin, detected in higher amounts relative to other phospholipids in Δ*lgt* -, has an important role in flagellar function [36], suggesting a direct link between lipoprotein modification and motility. Scanning electron microscopy (SEM) images show that the merodiploid *lgt* strain and Δ*lgt* + have a rougher cell surface compared to wild type (Fig. 5A). Lgt depleted cells are small, have spiky protrusions and lost cell polarity (Fig. 5A). Interestingly, the sheath surrounding the flagella filament was disintegrated in Δ*lgt*-as observed in transmission electron microscopy (TEM) images (Fig. 5B). Some sheaths were dislocated from the filament and in some cases were totally absent, leaving the filament naked for several micrometers. On some flagella, the sheath was agglomerating around or next to the flagellum forming vesicle-like structures. Cells in the revertant are rod shaped and flagella seem to possess an intact sheath similar to wild type (Fig. 5B), while the sheath defect was not completely restored in the Δ*lgt* suppressor although cells are rod shaped.

**Fig. 5.**
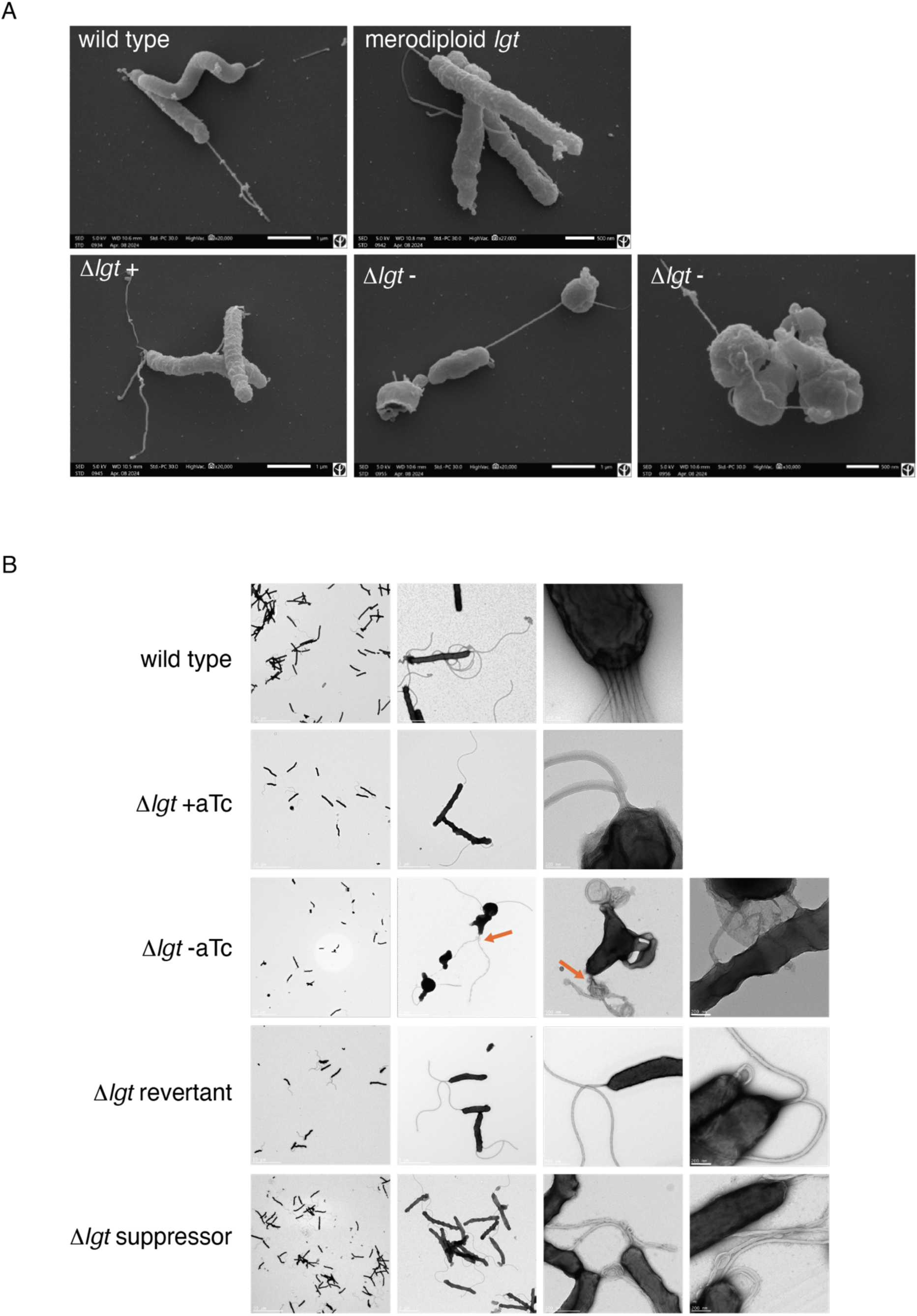
Lgt depletion affects morphology and flagellar sheath integrity. A) SEM images of wild type *H. pylori*, *lgt* merodiploid strain and Δ*lgt* with (+) and without (-) aTc. B) TEM images of wild type *H. pylori*, Δ*lgt* + *and* Δ*lgt* -, and revertant and suppressor. Strains were grown in BHI at 37°C under microaerophilic conditions for 24h (WT, Δ*lgt* +, Δ*lgt* - and Δ*lgt* suppressor) or 75h (Δ*lgt* revertant). Disintegrated flagella are indicated by arrows.

We next analyzed the capacity of the Δ*lgt* mutants to swim by counting the number of flagella and by performing motility plate assays (Fig. S6). *H. pylori* wild type strain B128, a highly motile strain [37], and *fli*A deletion mutants in N6 and B128 that are non-motile [38], were included as controls (Fig. 6A). The motility plate assay illustrates that *H. pylori* N6 is less motile than B128 (Fig. S6A and B). Δ*lgt* and the suppressor are non-motile, however, growth for 4 days on motility plates resulted in increased motility the of suppressor (Fig. S6A). While wild type *H. pylori* N6 contained up to eight polar flagella, Δ*lgt* + and suppressor possessed not more than two flagella on average per cell (Fig. S6C). These results reflect motility behavior of a cell population. Since our morphology analyses highlight the heterogenous character of the Δ*lgt* mutant, we explored live video microscopy to record motility of individual cells.

**Fig. 6.**
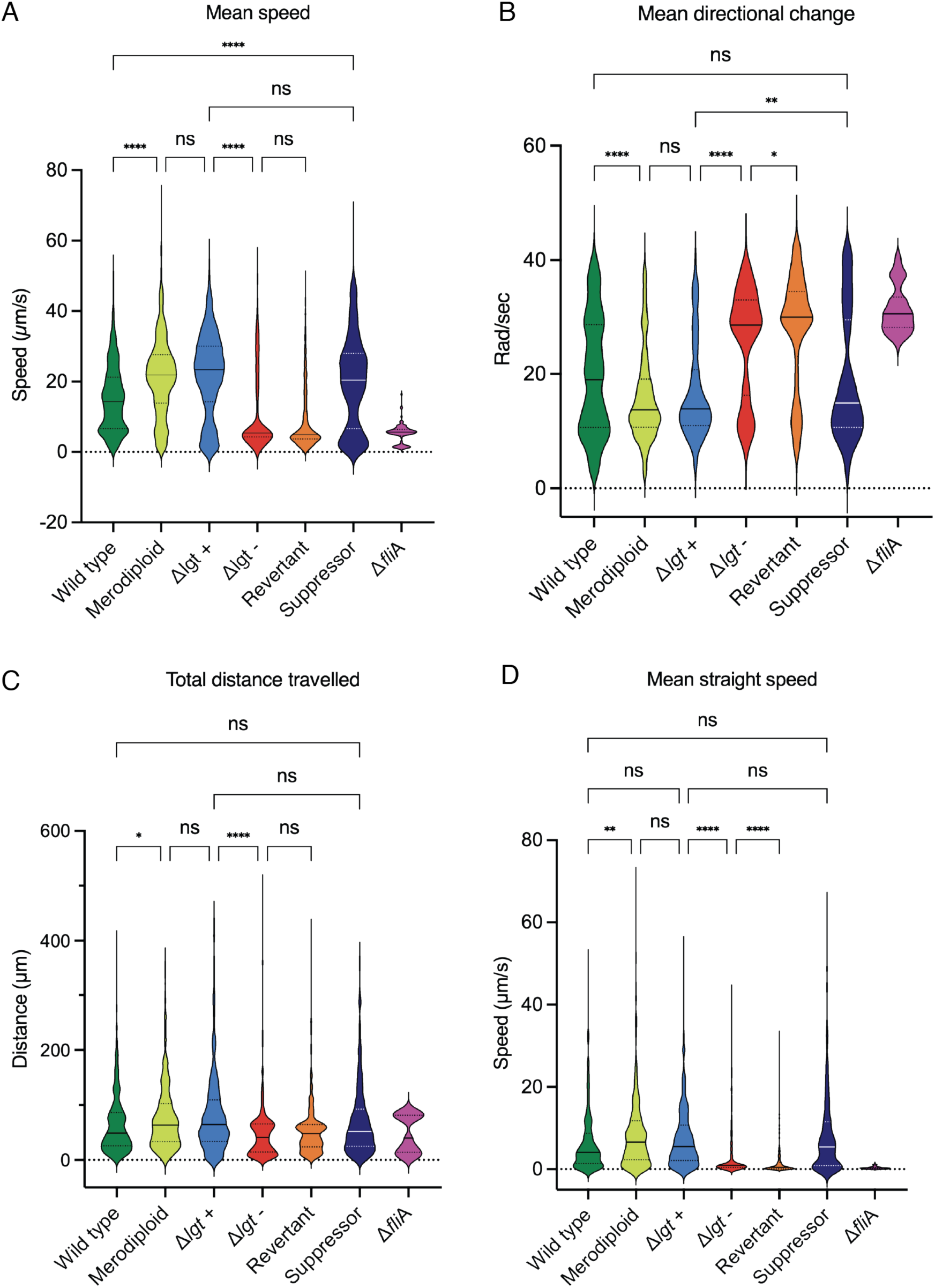
Depletion of Lgt leads to reduced motility and altered directional movement of *H. pylori*. A) Mean speed calculated by tracking 457 individual cells by video microscopy using the YOLO detection of Trackmate plugin in Fiji. B) Mean directional change calculated based on total angles between start- and endpoint of tracks. C) Total distance travelled presenting total of tracks of all cells per condition. D) Mean straight speed representing movement of cells in a straight line during the duration of the track. Mean values are indicated as solid lines, and one- and three-quarter values by dotted lines. Statistical analysis was performed by two-way ANOVA test with * p-value <0.05, ** p-value <0.01, *** p-value <0.001 and **** p-value <0.0001. Non-significant changes (p-value > 0.05) are not shown. Data were processed using Prism 10.4.1 (GraphPad).

The movement of cells was recorded by tracking cells in short videos in phase contrast mode with a 40-X objective on an epifluorescence microscope. Individual cells were automatically detected using the YOLO detector of the TrackMate plugin in Fiji [39]. The mean speed of cells in Δ*lgt* - and the revertant is lower than in Δ*lgt* +, while it increases to higher levels than wild type in the Δ*lgt* suppressor (Fig. 6A). The mean directional change indicates how cells move in time, *i*.*e*. cells that are non-motile and twist move locally around the start position, this is the case for Δ*fli*A (Fig. S6B). While cells in Δ*lgt* + behave like cells in the merodiploid *lgt* strain, both Δ*lgt* - and the Δ*lgt* revertant contain mainly cells that are immobile and some that are moving (mean directional change value is low) (Fig. 6B). Cells in Δ*lgt*- and the revertant move less far (Fig. 6C) and slower in a straight line than cells of Δ*lgt* +, while speed and directional movement was restored in the suppressor (Fig. 6C and D).

Overall, the results show that the heterogenous cell population in Δ*lgt* - and the revertant gives rise to a two-fold motility behavior and that motility is more efficient in the suppressor compared to the wild type.

## Discussion

*Helicobacter pylori* is a major risk for stomach cancer, for which antibiotic resistance is on the rise. Here we explored the role of Lgt on physiology and viability of *H. pylori* to obtain more insight into its potential as an antibacterial target. The absence of *lgt* expression led to a severe defect on growth and morphology leading to loss of viability and motility. *H. pylori* overcomes a defect in lipoprotein modification through transcriptional alterations in catabolism, energy metabolism, flagellar biogenesis, outer membrane protein production, phospholipid biosynthesis and DNA modification resulting in the appearance of revertants and suppressors. Only two lipoprotein encoding genes, uncharacterized HP_1440 and NucT (DNA nuclease involved in natural transformation), were directly affected while FapH (HP_0838) and HP_0018 were reported to have an important function in OM barrier conservation during flagellar rotation and a regulatory activity on peptidoglycan hydrolases, respectively. *H. pylori* lacks a classical bacterial SOS-response [40] but is able to resist oxidative stress. Peptidoglycan deacetylase (HP_0310) [41] and catalase (katA, HP_0875) [42] known to be responsive to oxidative stress were downregulated in the Δ*lgt* revertant. Downregulation of catabolism and energy metabolism may be a way to interrupt the normal cell cycle in order to survive. Furthermore, the stringent response may be activated where low levels of Lgt lead to low nutrient uptake via dedicated OM transporters and lack of this response may shut down stable RNA production [43] although a direct effect on *spo*T expression was not observed.

*H. pylori* has the capacity to regulate the expression of its gene arsenal when exposed to changing environments involving pH, nutrient availability and oxygen level through DNA modification by methylation and related processing by restriction enzyme systems. These changes often coincide with phase variation dependent gene expression where rearrangement of chromosomal regions results in an altered transcription profile [44]. The expression of flagellar genes is a well described example of phase variation regulated transcription. A *mod*H5 (HP_1522) deletion or *mod*H5 in the OFF state of phase variation results in less flagella and a decrease in motility [45]. Undermethylated regions in the chromosome of strain 26695 include virulence genes; *cag*PAI, *vac*A, *bab*A, *sab*A and *hop*Q and flagellar associated genes [46]. The *H. pylori* genome encodes twenty DNA methyltransferases (MTases) and for only a few a function has been assigned [47]. The expression of different MTase encoding genes were upregulated in the *lgt* revertant and one gene was downregulated in the suppressor. In the Lgt depletion strain the sole copy of *lgt* is under control of the *cag*UT promoter, which is regulated by methylation to control expression *cag*T (*cag*12) [48], which might explain the increase of *lgt* expression in the suppressor.

Several genes involved in flagellar biosynthesis were upregulated in the revertant, *i*.*e*. a chaperone (FlgN) together with filament-hook junction proteins (FlgL/K) it controls, ATP-ase complex/guide (FliJ), a flagellar export chaperone (FliS), a hook-length control protein (FliK), a MS ring component (FliF) and, finally, minor flagellin (FlaB). These are all structural components, no regulatory genes were affected, thus demonstrating that *H. pylori* compensates loss of lipoprotein modification through physical repair of flagella. The suppressor gained in swimming speed and motility even relative to the wild type strain. While few genes were upregulated: *fec*A, encoding an outer membrane iron dicitrate transport protein, HP_0057, encoding a Lpp20-like lipoprotein, *pab*B, encoding a para-benzoate synthetase and HP_0114. Except for HP_0114 (see below), a direct function in flagellar function is unclear. A combination of cell size and shape and flagellar rotation is likely to result in faster movement of cells, probably affecting infection and colonization capacity of *H. pylori*. Although loss of torque (2 versus 7 flagella) might prevent the bacterium to penetrate the mucus layer.

We observed important changes in the phospholipid composition of membranes when lipoprotein modification was disturbed through reduced expression of *lgt* causing an effect on cell envelope architecture. The fatty acid composition of two lipoproteins of *H. pylori* has been reported; both CagT and Lpp20 contained C16:0 and C18:0. We reported that Lgt of *H. pylori* is functional in *E. coli* [16] in which lipoproteins at least contain palmitate (C16:0) [49] and PG is the preferred acyl donor [17]. A minor PG species (PG34:1) in *H. pylori* contains C16:0/C18:1 fatty acids and could possibly function as substrate in the Lgt reaction. Future studies on substrate specificity of Lgt enzymes are required to pinpoint potential differences in lipoprotein modification in bacterial pathogens. The pathway for phosphatidylglycerol and cardiolipin synthesis involves several enzymes starting from phosphatidic acid (PtdOH) (Fig. S2). First, cytidine diphosphate-diacylglycerol (CDP-DAG) is formed through catalysis by CdsA. PgsA catalyzes the conversion of CDP-DAG and glycerol-3-phosphate (G3P) to phosphatidylglycerophosphate (PGP) followed by dephosphorylation by PgpA resulting in PG. Cardiolipin synthase (ClsC; HP_0190) is the major cardiolipin synthase in *H. pylori* that combines two molecules of PG and loss of this enzyme blocks flagella assembly [36]. ClsC localizes to the cell pole, an area with negative curvature and the location of flagella [29]. It is known that CL stabilizes membrane proteins and complexes ranging from bacteria to mitochondria [50,51]. CL is still detected in *H. pylori* membranes in absence of ClsC, suggesting that additional cardiolipin synthase(s) exist [36]. Lack of lipoprotein maturation due to depletion of Lgt leads to overproduction of CL. ClsC is a predicted lipoprotein [52], suggesting that the enzyme exists in a non-lipidated state under these conditions. This raises several interesting questions about the correlation between the polar localization of ClsC and its role in flagellar integrity. A recent report demonstrated that ClsO (predicted lipoprotein), a ClsC homolog in *Acinetobacter baumannii*, is a surface exposed cardiolipin synthase [52]. Cardiolipin is a negatively charged phospholipid and both major and minor flagellins FlaA and FlaB are post-translationally modified with seven and nine glycans, respectively, resulting in negatively charged filaments. This could explain the disorganization of the sheath through repulsive interaction between CL and FlaAB, leading to sheath break, which is restored in the revertant by higher expression of *fla*B, lower levels of CL and an increase in lysophospholipids. A specific role in this process could be attributed to HP_0114, a motility associated glycosyltransferase, known to decorate FlaAB with glycans and which expression is upregulated in both the revertant and the suppressor. This protein is important for motility since a HP_0114 deletion mutant is non-flagellated [53]. While the expression level of two genes involved in phospholipid biosynthesis, *cdh* and *pgs*A, was altered no genes encoding phospholipases were identified to explain the increase in lysophospholipids. The many uncharacterized and unknown genes of which expression was affected in absence of *lgt* expression indicates that many regulatory mechanisms are to be discovered.

In conclusion, incomplete lipoprotein modification in *H. pylori* results in short- and long-term adaptation of gene expression that affect cell envelope integrity and related OMP and flagellar function. Our findings are insightful for novel antibacterial strategies where lipoprotein modification enzymes would be a promising target to be explored possibly in synergy with existing antibiotics.

## Acknowledgements

Financial aid was obtained by a grant from the “Programmes Transversaux de Recherche” (PTR 626-23) from Institut Pasteur, by the Labex “Integrative Biology of Emerging Infectious Diseases” (ANR-10-LBX-62 IBEID) and by the Fondation pour la Recherche Medicale (FRM) (EQU202403018034). TR received a PhD fellowship from the French Ministry of Research and Education. We thank M. Monot and L. Lemée (Biomics Platform, C2RT, Institut Pasteur) for RNAseq analysis financed by France Génomique (ANR-10-INBS-09) and IBISA. Etienne Kornobis (Biomics Platform) and Damien Mornico (Bioinformatics HUB, Institut Pasteur) for whole genome Illumina sequence analysis. We thank S. Phan and J-Y Tinevez for help and suggestions on video microscopy analysis and C. Schmidt and C. Scandola for training and assistance with electron microscopy. We are grateful to M. McClain and T. Clover for providing chromosomal DNA from strain VM176. We thank all members of the BGPB Unit for their support and constructive discussions.

## Material and Methods

### Bacterial growth media and culture conditions

*H. pylori* strains (Table S1) were grown on Blood (BA) or Brucella Broth (BBA) agar plates, or in liquid Brain Heart Infusion (BHI) medium containing an antibiotic-antifungal mix (0.3 μg/mL polymyxin B, 2.5 μg/mL amphotericin B, 12.5 μg/mL vancomycin, 6.25 µg/mL trimethoprim) under shaking and microaerophilic conditions (6% O_2_, 10% CO_2_). Antibiotics kanamycin (Kan 20 μg/mL), chloramphenicol (Cm 4 μg/mL) and gentamycin (Gm 5 μg/mL) were added when required. BA plates were complemented with 10% defibrinated horse blood. BBA plates and BHI cultures were supplemented with 10% of decomplemented fetal calf serum (FCS). Anhydrotetracycline (aTc; 100 ng/mL) was added to the merodiploid *lgt* and Δ*lgt* + strains in order to express *lgt* from the *cagUT* promoter. Growth in liquid medium was either performed in cell culture flasks (25 cm^2^ for 10 mL, 75 cm^2^ for 50 mL and 150 cm^2^ 100 mL cultures) or in 24-well plates (1 mL cultures). For microscopy, RNAseq and phospholipid analyses, time samples were taken according to the growth curve shown in Fig. S1. The time point of exponential growth of the Δ*lgt* revertant is the same for cultures in 24-well plate and 25 cm^2^ flasks. Growth curves were obtained by measuring OD_600_ in a Spark Plate Reader (TECAN) or a cuvette spectrophotometer (Eppendorf). For growth in the TECAN, bacteria were inoculated in medium and incubated in the presence of an adapted humidity cassette. Atmosphere and temperature were controlled, and incubation was performed under orbital shaking at 216 rpm and 2.5 mm of amplitude. Data were analyzed using Prism 10.4.1 (GraphPad).

### Construction of bacterial strains

A Lgt depletion strain was constructed in *H. pylori* N6 following a 3-step procedure. Chromosomal DNA from strain VM176 [14] was used to amplify *ure*A-*tet*R and *hp_0630*-*lgt*-*hp_0631*. The PCR fragments were sequentially transformed into N6 and selected for growth on solid medium containing Kan and Cm, respectively, resulting in merodiploid strain NBHP2 (Table S1). The endogenous *lgt* gene (*hp_0955*) was replaced by a Gm^R^ cassette from plasmid pUC18-Gm through transformation of a PCR fragment from a 3-steps PCR process using primer pair 1 and 2 and primer pair 3 and 4 (Table S1). All constructs were verified by sequencing of PCR products of inserted loci (Eurofins or Plasmidsaurus).

### Determination of cell viability

The OD_600_ of bacterial cultures was normalized to 1.0. Serial dilutions were performed to a final dilution of 10^-7^. Spots of 2.5 µL were spotted on BBA plates containing 40 μg/mL triphenyl tetrazolium chloride (TTC) and complemented with aTc when required. Plates were incubated at 37°C under microaerophilic conditions for four days. The number of colony forming units (CFU) was determined by counting colonies from images of plates taken on a ChemiDoc Imager (BioRad). Graphs were performed using Prism 10.4.1 (GraphPad).

### Transcriptome analysis

Total RNA was isolated from 4 independent replicates of *H. pylori* cultures grown in 10 mL BHI for 24h or 75h (Δ*lgt* revertant) (see Fig. S1B) using a NucleoSpin RNA extraction kit (Macherey-Nagel) following the manufacturer’s protocol. Quantity and quality measurements were performed using Qubit Fluorometer (Thermo Fischer Scientific) and the Agilent 2100 Bioanalyzer (Agilent Technologies). Sequencing libraries were constructed using an Illumina Stranded Total RNA prep, Ligation with Ribo-Zero Plus kit (Illumina, USA) with custom primer design on rRNA following the supplier’s recommendations. The efficiency of rRNA depletion was evaluated by shallow sequencing. RNA sequencing was performed on an Illumina NextSeq 2000 platform for a target of 10M reads/sample. The RNAseq analysis was performed with Sequana 0.15.3 using the RNAseq pipeline 0.16 built on top of Snakemake 7.32/4. Reads were trimmed from adapters using Fastp 0.22.0 then mapped to the *H. pylori* genome (RefSeq AE000511.1) using Bowtie2 and STAR. Quality control statistics were summarized using MultiQC 1.11. Statistical analysis on the count matrix was performed to identify differentially regulated genes. Clustering of transcriptomic profiles were assessed using a Principal Component Analysis (PCA). Differential expression testing was conducted using DESeq2 library 1.34.0 scripts indicating the significance (Benjamini-Hochberg adjusted p-values, false discovery rate <0.05) and the effect size (fold change) for each comparison. Genes of N6 were annotated using the genome of *H. pylori* strain 26695 and manually added as gene ontology (GO) annotation for predicted protein function when annotations were not found.

### Whole genome sequencing

DNA was isolated from 10 mL *H. pylori* cultures grown in BHI for 24h or 75h (Δ*lgt* revertant). Samples were fragmented with a Covaris M220 focused ultrasonicator (Covaris Ltd., Brighton, United Kingdom) using microTUBE-15 to 350 bp. The TruSeq DNA PCR-free libraries prep kit (Illumina) was used as instructed by the manufacturer. Sequencing was performed on an Illumina ISeq 100 platform. Raw reads were processed with an in-house bioinformatics pipeline for quality control. We next used the variant calling pipeline (v0.11.0), which is available online (https://github.com/sequana/variant_calling) to identify mutations. Analysis of variants was further performed using the polymorphism mode in Breseq version 0.35.5 [54] on the reference genome of N6. Nanopore sequencing (Plasmidsaurus) was performed on chromosomal DNA isolated from three independent Δ*lgt* suppressor strains.

### Scanning Electron Microscopy

Liquid precultures of *H. pylori* strains were inoculated in 1 mL BHI containing 10% FCS at a starting OD_600_ of 0.05, with or without aTc, in a 24-well plate. Bacteria were harvested at exponential phase (24h) or 75h (Δ*lgt* revertant) and fixed in growth medium with 2.5% glutaraldehyde for 15 minutes at room temperature. A filter was added to each well and cells were left to sediment during 1 h at room temperature to preserve the flagella. Bacteria were then postfixed for 1 h 30 min in 1% osmium tetroxide in 0.1 M HEPES buffer (pH 7.2), followed by rinsing once with distilled water. Samples were dehydrated through a graded series of 25, 50, 75, 95, and 100% ethanol solution, followed by critical point drying with CO_2_. Dried specimens were sputtered with 20-nm gold palladium with an ion beam coater and examined and photographed with a JEOL IT 700 HR field emission SEM operating at 5 kV. Images were acquired with the upper secondary electron (SE) detector in-lens upper secondary electron detector (SEI). Images were processed with Adobe Photoshop for data presentation.

### Transmission Electron Microscopy

Liquid precultures of *H. pylori* strains were inoculated in 10 mL of BHI containing 10% FCS at a starting OD_600_ of 0.05, with or without aTc, in 25cm^3^ flasks. Bacteria were fixed in growth medium at exponential phase (24h) or 75h (Δ*lgt* revertant) with 2.5% glutaraldehyde for 15 minutes at room temperature and were then harvested through centrifugation at 800 x g to preserve the flagella, washed three times in 20 mM HEPES buffer (pH 7.2) and finally resuspended in 20 mM HEPES buffer (pH 7.2) at an OD_600_ of 0.1. The cell suspension (10 μL) was absorbed to 200-mesh, Formvar carbon-coated-copper glow discharged grids (Electron Microscopy Sciences UK), and incubated for 20 minutes at room temperature. Grids were washed three times by letting them incubate in a drop of water for two minutes to remove the excess of non-adhered cells. Samples were stained with 2% aqueous uranyl acetate for 10 seconds. Filter paper was used to dry the grids. Images were taken with a transmission electron microscope Tecnai Biotwin T12 (FEI-Thermo Fisher Company) at 120kV accelerating voltage with a Gatan RIO 16 4k camera. Images were processed with Adobe Photoshop (v.27.9.1) for data presentation.

### Phospholipid and fatty acid analyses

The glycerolipids were extracted from membranes fractions that were prepared from 50 mL *H. pylori* cell cultures grown in BHI to mid-logarithmic phase. Cell pellets were harvested by centrifugation at 4,000 x g for 15 min at 4°C and resuspended in 20 mM HEPES buffer (pH 7.4). Cells were broken by two passages in a French Press at 10 psi and unbroken cells and debris were removed by centrifugation at 10,000 x g for 10 min at 4°C. Membrane fractions were collected by centrifugation at 100,000 x g for 60 min at 4°C and resuspended in 20 mM HEPES buffer (pH 7.4) to 0.4 OD_600_ units/μL. One milliliter of methanol preheated to 85°C was added to 50 μL sample and incubated at 85°C for 5 min. Subsequently, 3 mL of methyl-*tert*-butyl ether (MTBE) was added, and the mixture was incubated at room temperature with shaking for 15 min. Phase separation was induced by adding 1 mL of water. After centrifugation at 3000 × *g* for 5 min, the upper organic phase was collected and evaporated under a stream of nitrogen. The dried lipid extracts were resuspended in 500 µl of a mixture acetonitrile/isopropanol/water/ammonium formiate 10mM (65/35/4/1, v/v/v/v) and stored at −20 °C.

For fatty acid composition analysis, 250 µl of the extracted lipids was dried, converted to fatty acid methyl esters (FAMEs) by acid-catalyzed transmethylation and analyzed by gas chromatography coupled to flame ionization detection and mass spectrometry (GC-FID-MS). Briefly 1 ml of a solution containing methanol with 5% (v/v) sulfuric acid was added to dried extract in sealed glass tubes. Samples were incubated at 85°C for 90 min. After cooling down, FAMEs and hydrocarbons were extracted by adding 500 µL hexane and 1.5 ml 0.9% (w/v) NaCl. Samples were mixed and the organic phase was separated from the aqueous phase by centrifugation at 3,000g for 2 min. The hexane phase was recovered and 1 µL was injected in the GC-MS/FID. GC-MS/FID analyses were carried out on an Agilent 7890A gas chromatographer coupled to an Agilent 5975C mass spectrometer (simple quadrupole). A Zebron 7HG-G007-11 (Phenomenex) polar capillary column (length 30 m, internal diameter 0.25 mm, and film thickness 0.25 µm) was used. Helium carrier gas was at 1 mL min−1. Oven temperature was programmed with an initial 2-min hold time at 60°C, a first ramp from 60 to 150°C at 20°C⋅min−1, then a second ramp from 150 to 240°C at 6°C⋅min−1 and a final 3-min hold time at 240°C. Samples were injected in splitless mode (1 min) at 250°C. The MS was run in full scan over 40 to 350 amu (electron impact ionization at 70 eV). FAMEs were identified based on mass spectrum and quantified using FID trace.

Lipidomic analysis was performed using a Thermo Vanquish UPLC system coupled with a Thermo Orbitrap Q Exactive mass spectrometer operated in data-dependent acquisition (DDA) mode. A 2µL aliquot of each sample was injected onto a Kinetex XB-C18 column (150 mm length, 2.1 mm internal diameter, 1.7 μm particle size) at a flow rate of 0.3 mL/min, with the column temperature maintained at 45 °C. The mobile phase consisted of solvent A (acetonitrile/water 60:40, *v*/*v,* containing 10 mM ammonium acetate) and solvent B (isopropanol/acetonitrile 90:10, *v*/*v,* containing 10 mM ammonium acetate). Lipids were separated using the following gradient of solvent B: the gradient started at 27% B for 0.5 min, increased linearly to 97% B over the next 24 minutes, was held constant during 2 min, returned linearly to 27% B and was held constant until the end of the run (32 min). The Q-Exactive mass spectrometer was used in negative ion mode with the following source parameters: sheath gas at 50 arbitrary units (AU); sweep gas at 2 AU; auxiliary gas at 15 AU; spray voltage at 2.5 kV; capillary temperature, 300 °C; S-lens RF level, 50; scan range, *m*/*z* 200–1100. MS/MS spectra were acquired using automatic gain control target and stepped normalized collision energy of 25 eV, 30 eV and 40 eV. The acquired raw data were processed using TraceFinder (Thermo Fisher Scientific, Waltham, MA, USA). Lipid molecular species were identified based on accurate mass measurements and lipid fragmentation patterns.

Standard commercial phospholipids used were; 1,2-dipalmitoyl-sn-glycerol-3-phospho-(1’-rac-glycerol) (phosphatidylglycerol, PG), 1,2-dipalmitoyl-*sn*-glycerol-3-phosphoethanolamine (phosphatidylethanolamine, PE), lysophosphatidylethanolamine (LPE) 1,3-bis[1-palmitoyl-2-oleolyl-*sn*-glycero-3-phospho]-glycerol (cardiolipin, CL), phosphatidylcholine (PC), lysophosphatidylcholine (LPC). The amount of phospholipid was calculated by peak area from UPLC-MS correlated to the total amount of FAMEs calculated from FID traces in the sample. Data were analyzed with Prism 10.4.1 (GraphPad).

### Epifluorescence microscopy imaging and video microscopy

For morphological analyses, *H. pylori* cells were fixed in 2.5% glutaraldehyde in growth medium for 15 minutes at room temperature and washed three times in 20 mM HEPES buffer (pH 7.4). DNA was stained with Hoechst 2333 (10 μg/mL, Biotium) and membranes with FM4-64X (0.2 μg/mL, Invitrogen) for 15 minutes at room temperature in the dark. Cells were washed three times in 20 mM HEPES buffer (pH 7.4) with centrifugation at 10,000 x g for 1 min. Cells were immobilized on agar pads (1% agarose in H_2_O) and imaged with a Zeiss Axio Observer inverted epi-fluorescence microscope. Images were analyzed using the MicrobeJ version 5.13I plugin of Fiji [55]. Cell parameters (length and width) were measured and collected automatically. Representative images were processed using Adobe Photoshop 27.9.1and Adobe Illustrator 30.8.0.

Short videos (15 sec) of live *H. pylori* cells were recorded using a 6-channel μ-Slide VI 0.4 μ-Bioinert (IBIDI) by phase contrast microscopy with a 40-X objective under microaerophilic conditions at 37°C. Individual cells were automatically detected using the YOLO detector of the TrackMate plugin [39] in Fiji, and tracks drawn with the Simple-LAP Tracker [56] and further analyzed for motility parameters following the developers’ protocol [39].

### Cell motility analysis

A motility plate assay was used to determine motility of a total cell population. *H. pylori* cells were collected from BA plates to inoculate precultures in BHI. After 16h of growth at 37°C under microaerophilic conditions. 2 µL of culture (normalized to OD_600_ of 1.0) was spotted onto BBA plates containing 0.35% agar and 40 μg/mL TTC. Plates were incubated at 37°C under microaerophilic conditions for four days. The diameter of the red halo was measured for each strain from images taken on a ChemiDoc Imager (BioRad).

---

**Suppl. Fig. 1.**
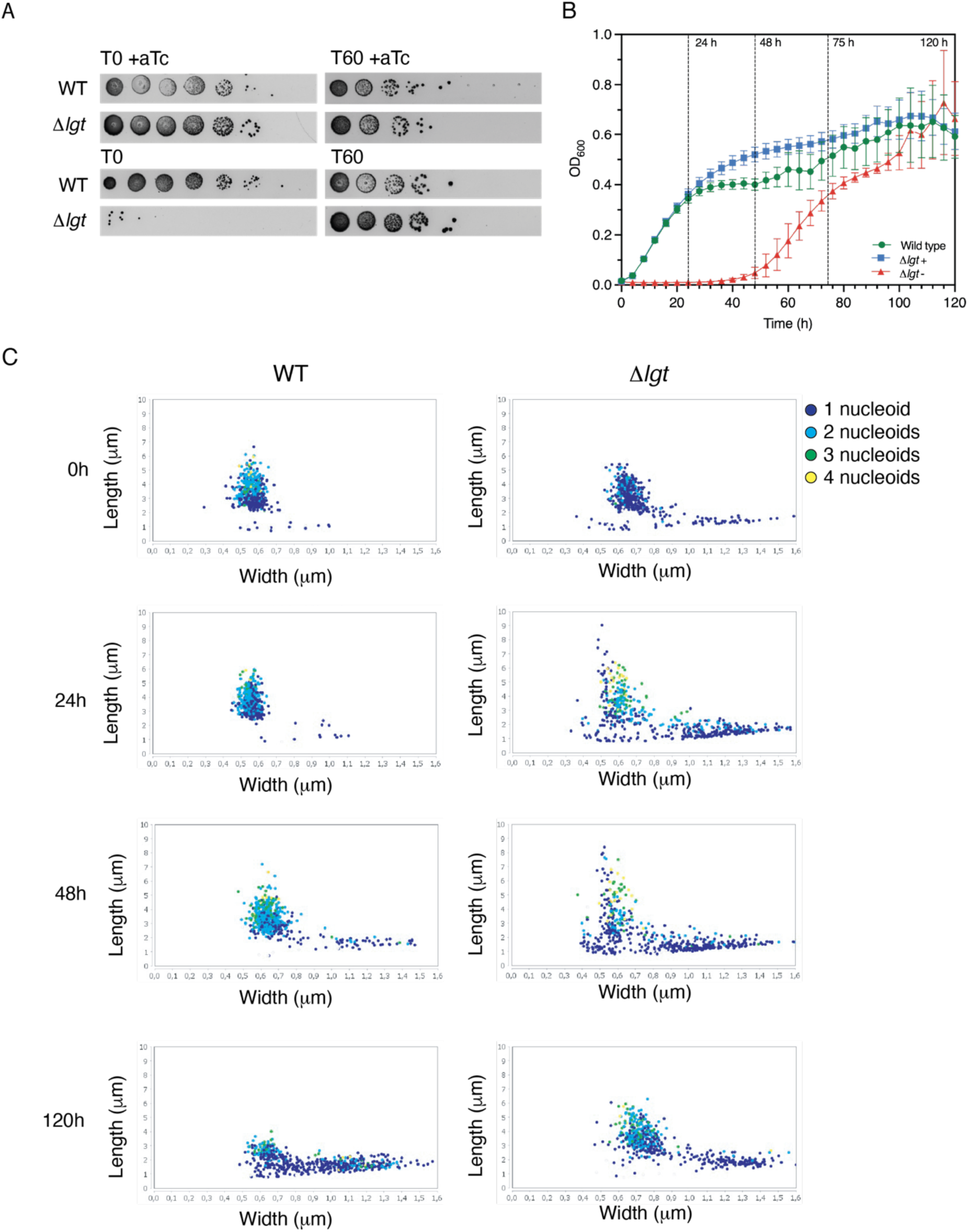
A. Spot dilution assay corresponding to data presented in Fig. 1C. B. Growth curve of wild type and Δ*lgt* strain in 1 mL cultures in 24-well plate. Time of onset of revertant growth corresponds with flask grown cultures of 10 mL. C. Evolution of cell shape in wild type and Δ*lgt* (-aTc) during growth over 120h. Dots represents single cells with one nucleoid (dark blue), two nucleoids (light blue), three nucleoids (green) or four nucleoids (yellow). Quantification of cell parameters was performed using the MicrobeJ plugin in Fiji with 500-1000 cells per strain and per condition. Data were processed using Prism 10.4.1 (GraphPad).

**Suppl. Fig. 2.**
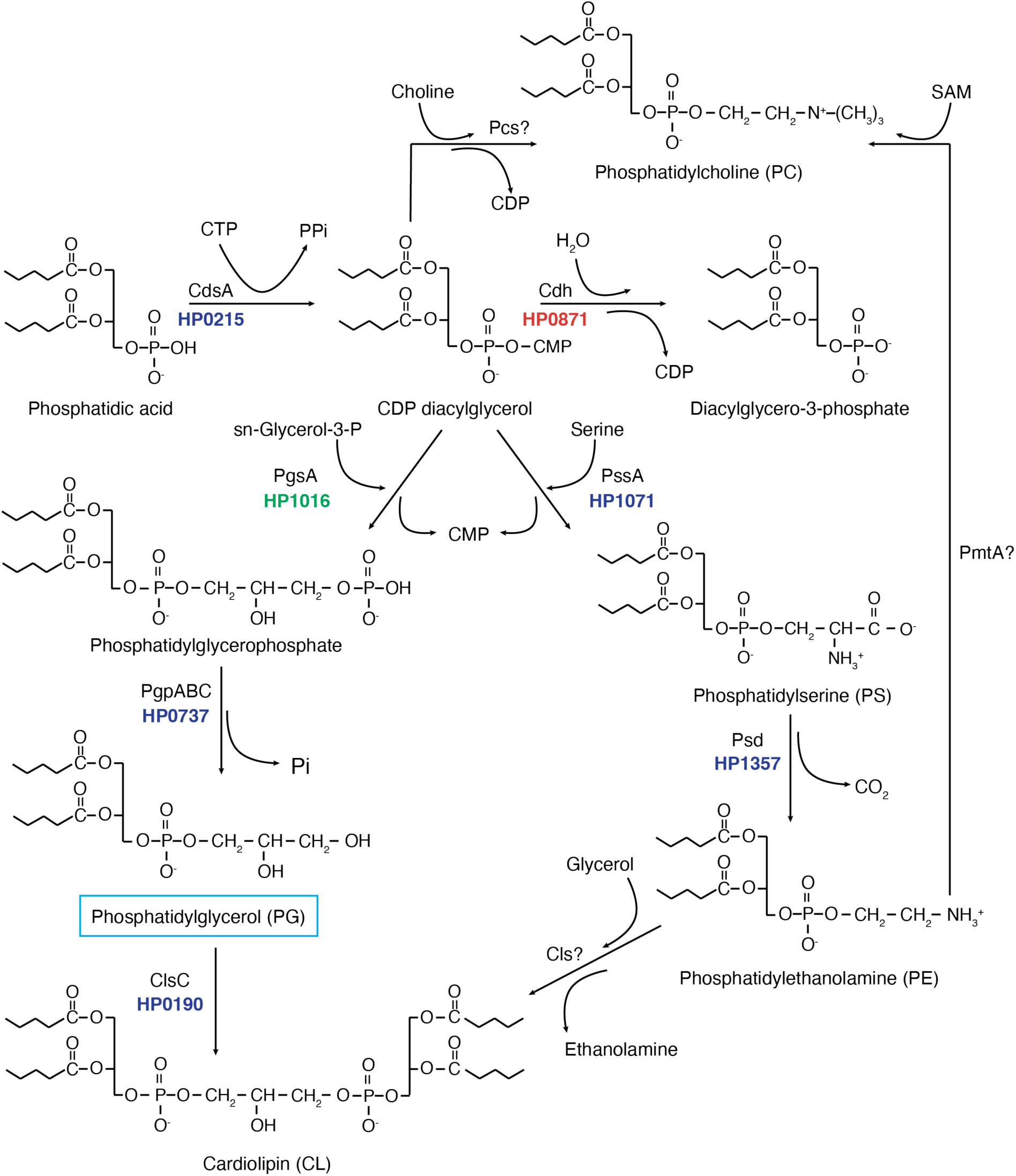
Phospholipid synthesis is altered to compensate for loss of Lgt function in *H. pylori*. Phospholipid biosynthesis pathway based on *E. coli* [57] and phosphatidylcholine synthesis postulated according to the pathway described for various bacterial species [58]. HP_1016 (*pgsA*) is upregulated and HP_0871 (*cdh*) is downregulated in the Δ*lgt* revertant, while other synthetic genes are not affected. *Cls*C encodes the main cardiolipin synthase in *H. pylori* and homologous genes of *cslA*/*B* involved in the synthesis of CL from PE have not been identified. CDP-DAG synthase (CdsA), CDP-diacylglycerol pyrophosphatase (Cdh), phosphatidylcholine synthase (Pcs), phosphatidylglycerophosphate synthase (PgsA), phosphatidylserine decarboxylase (Psd), phosphatidylserine synthase (PssA), phosphatidylglycerophosphate phosphatases (PgpABC), phosphatidylethanolamine N-methyltransferase (PmtA), *S*-adenosyl-L-methionine (SAM).

**Suppl. Fig. 3.**
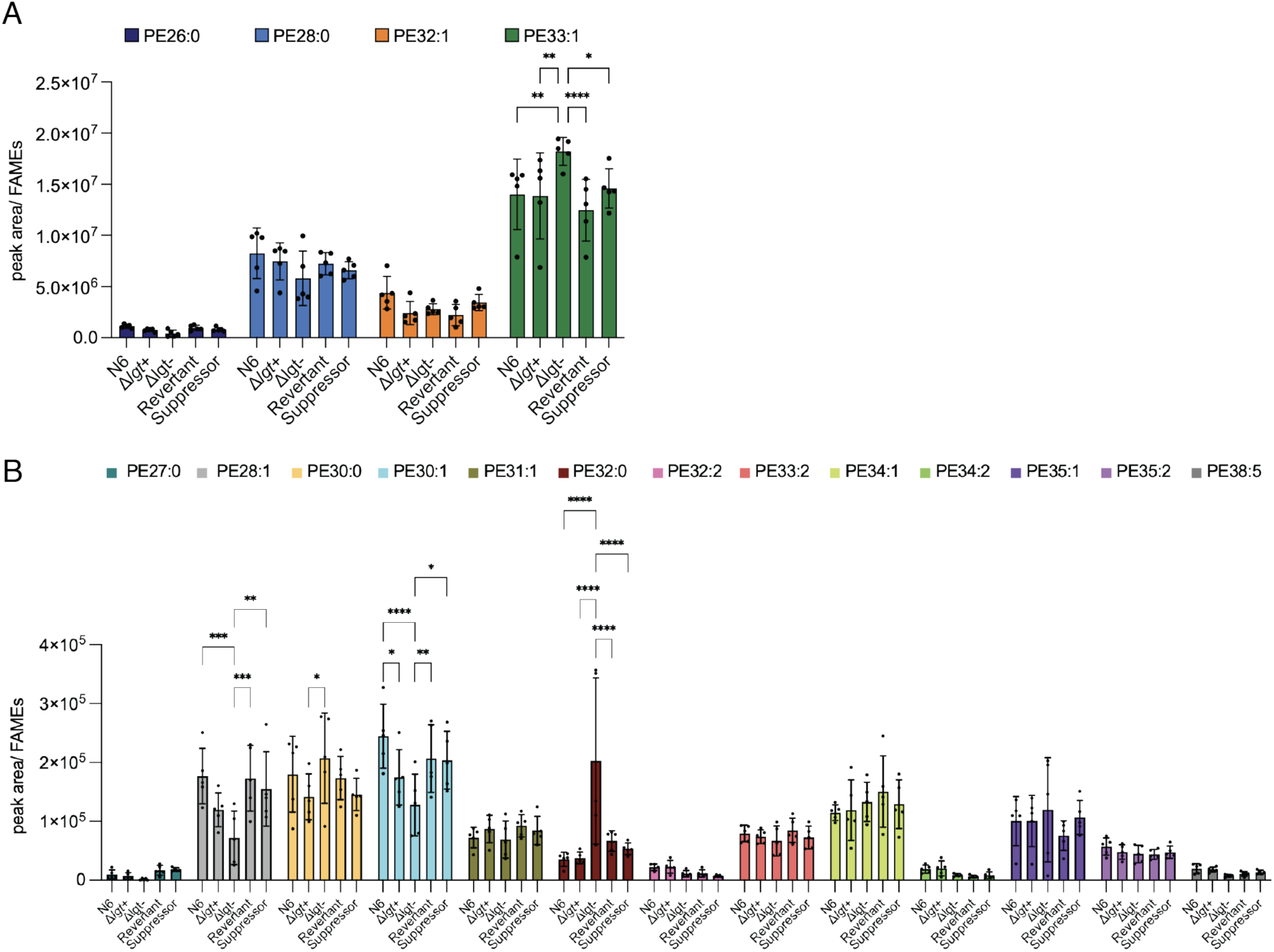
A) Quantification of major PE species relative to the total amount of fatty acids. E) Quantification of minor PE species relative to the total amount of fatty acids. Analysis was performed with data from five biological replicates. Statistical analysis was performed by two-way ANOVA test with * p-value <0.05, ** p-value <0.01, *** p-value <0.001 and **** p-value <0.0001. Non-significant (p-value >0.05) differences are not indicated. Data were processed using Prism 10.4.1 (GraphPad).

**Suppl. Fig. 4.**
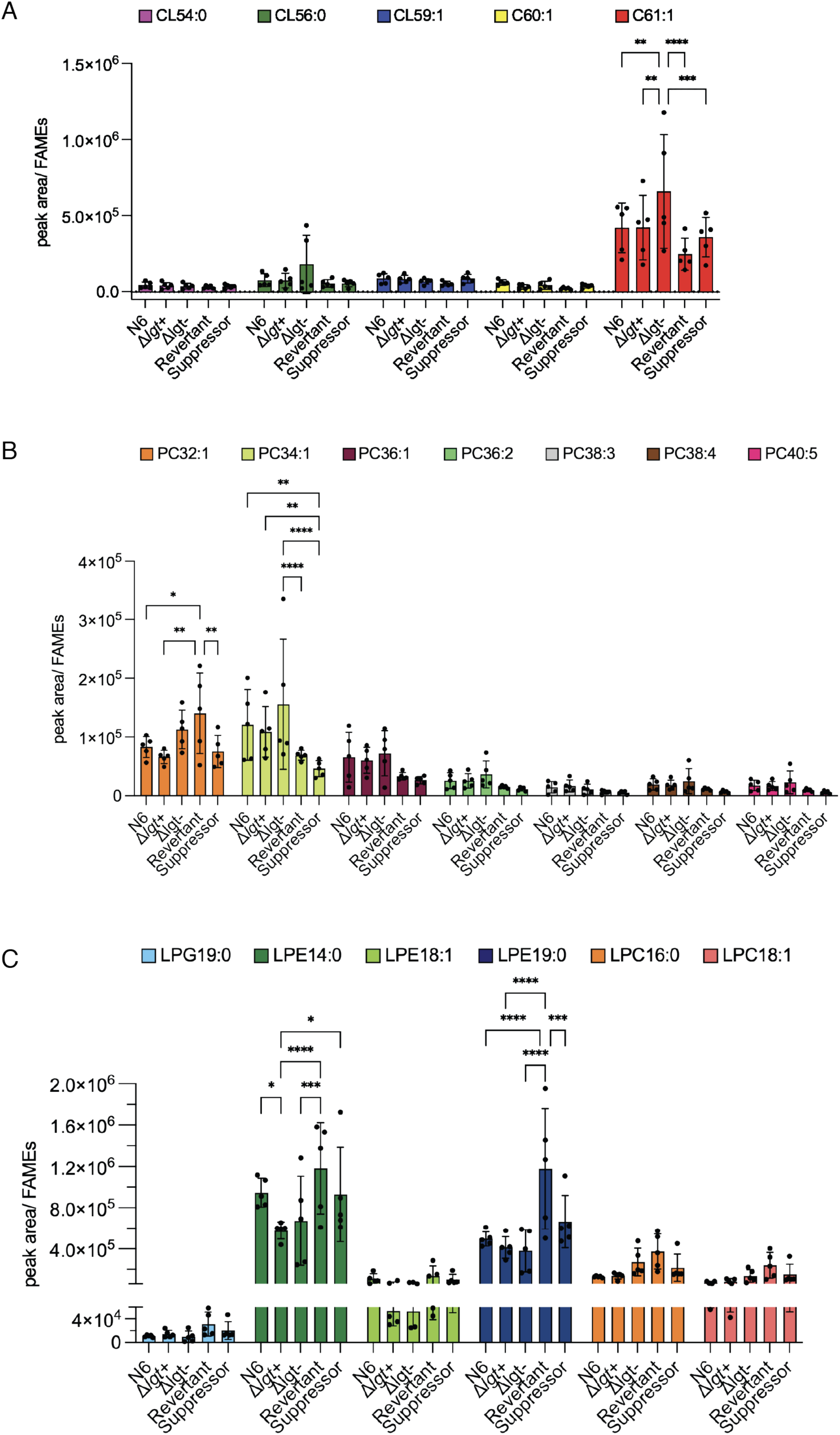
A) Quantification of CL species relative to the total amount of fatty acids. B) Quantification of PC relative to the total amount of fatty acids. D) Quantification of lysophospholipids relative to the total amount of fatty acids. Statistical analysis was performed by two-way ANOVA test with * p-value <0.05, ** p-value <0.01, *** p-value <0.001 and **** p-value <0.0001. Non-significant (p-value >0.05) differences are not indicated. Data were processed using Prism 10.4.1 (GraphPad).

**Suppl. Fig. 5.**
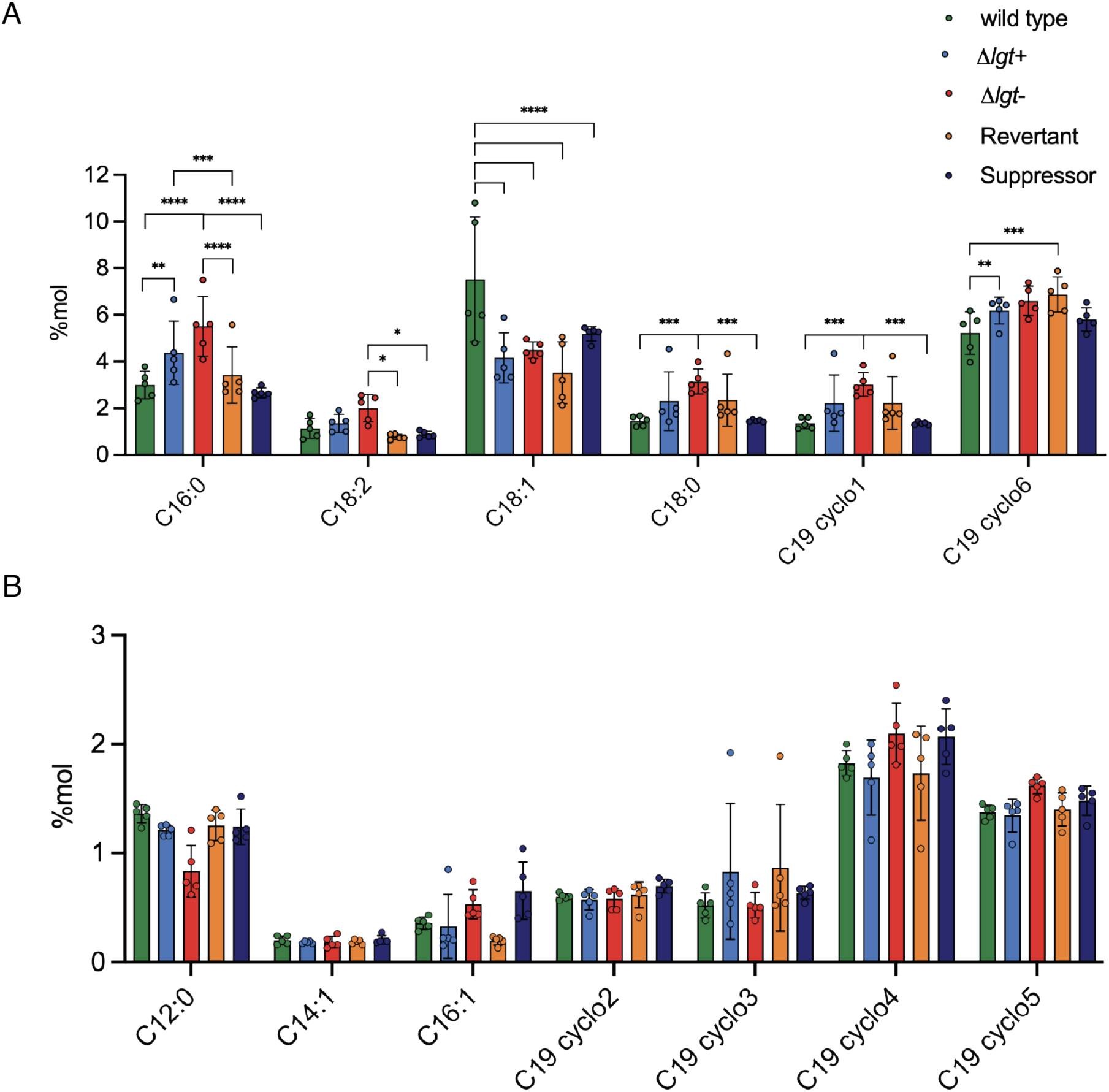
A) Quantification of fatty acid moiety of membrane phospholipids as percentage of total fatty acids in the sample. Relative amount of major fatty acid C14:0 is not shown. Significant changes between fatty acid species are indicated as analyzed by two-way ANOVA statistical analyses with * p-value <0.05, ** p-value <0.01, *** p-value <0.001 and **** p-value <0.0001. B) Non-significant changes (p-value >0.05) were observed with minor fatty acid species of phospholipids in *H. pylori* membranes. Data were processed using Prism 10.4.1 (GraphPad).

**Suppl. Fig. 6.**
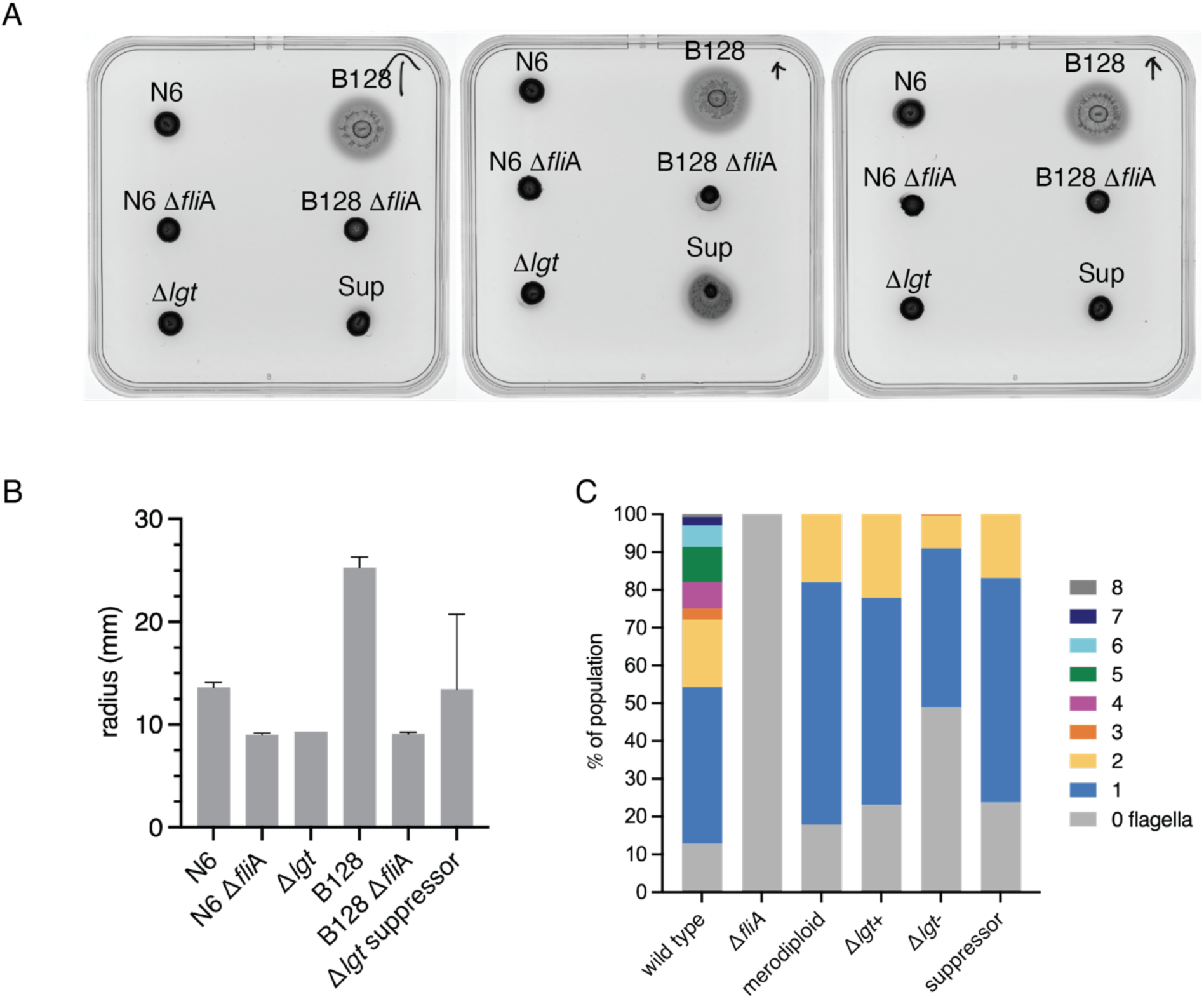
A) Plate motility assay with *H. pylori* wild type strains N6 and B128, with corresponding *fli*A deletion mutants and Δ*lgt*. Cells were spotted onto motility agar plates and grown at 37°C under microaerophilic conditions for 4 days. B) Motility was quantified by measuring the diameter of each halo from images taken from plates shown in panel A. The experiment was performed in triplicate. C) Number of flagella was determined from 100 cells in each strain and condition in TEM images.

## Supplementary Tables

**Suppl. Table 1.**
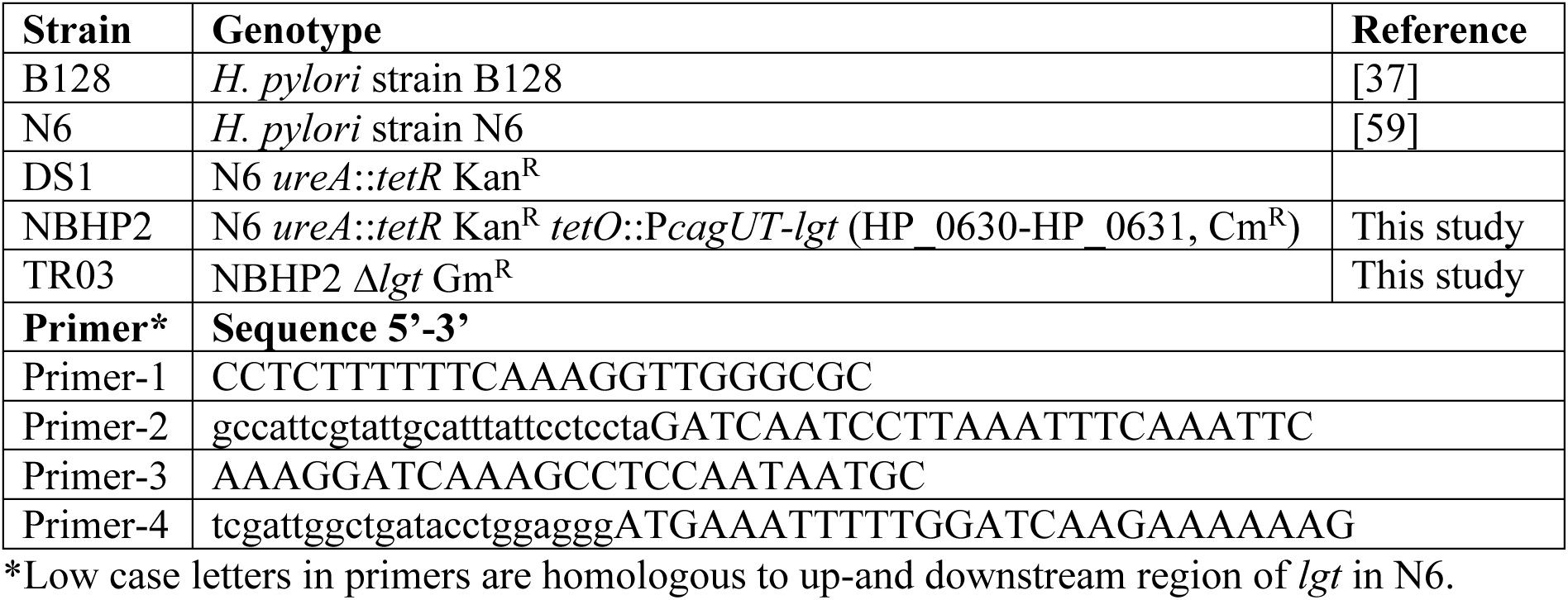
Strains and primers used in this study.

**Suppl. Table 2.**
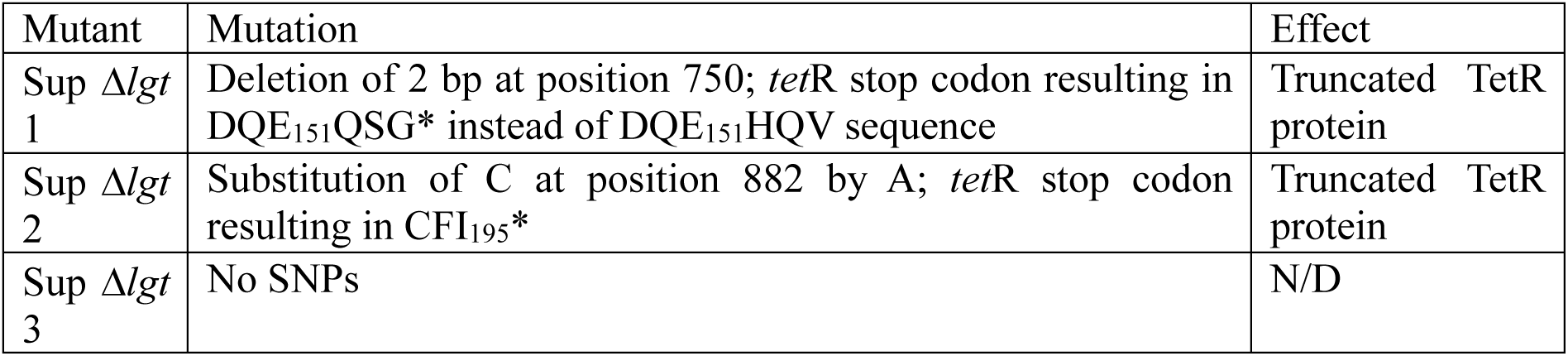
Mutations identified by WGS analysis in Δ*lgt* suppressor mutants.

| Mutant | Mutation | Effect |
| --- | --- | --- |
| Sup Δ*lgt* 1 | Deletion of 2 bp at position 750; *tetR* stop codon resulting in DQE<sub>151</sub>QSG* instead of DQE<sub>151</sub>HQV sequence | Truncated TetR protein |
| Sup Δ*lgt* 2 | Substitution of C at position 882 by A; *tetR* stop codon resulting in CFI<sub>195</sub>* | Truncated TetR protein |
| Sup Δ*lgt* 3 | No SNPs | N/D |

**Suppl. Table 3.** Level of *lgt* expression.

| Strain 1 | Strain 2 | Fold Change |
| --- | --- | --- |
| Wild type | Merodiploid | 0.35 |
| Wild type | Δ*lgt*+ | 1.90 |
| Wild type | Revertant | 3.15 |
| Wild type | Suppressor | 0.42 |
| Merodiploid | Suppressor | 0.88 |
| Δ*lgt*+ | Merodiploid | 0.15 |
| Δ*lgt*+ | Suppressor | 0.13 |
| Revertant | Merodiploid | 0.22 |
| Revertant | Δ*lgt*+ | 1.50 |
| Revertant | Suppressor | 0.20 |

**Suppl. Table 4.** All RNA seq data. See appendix.

**Suppl. Table 5.**
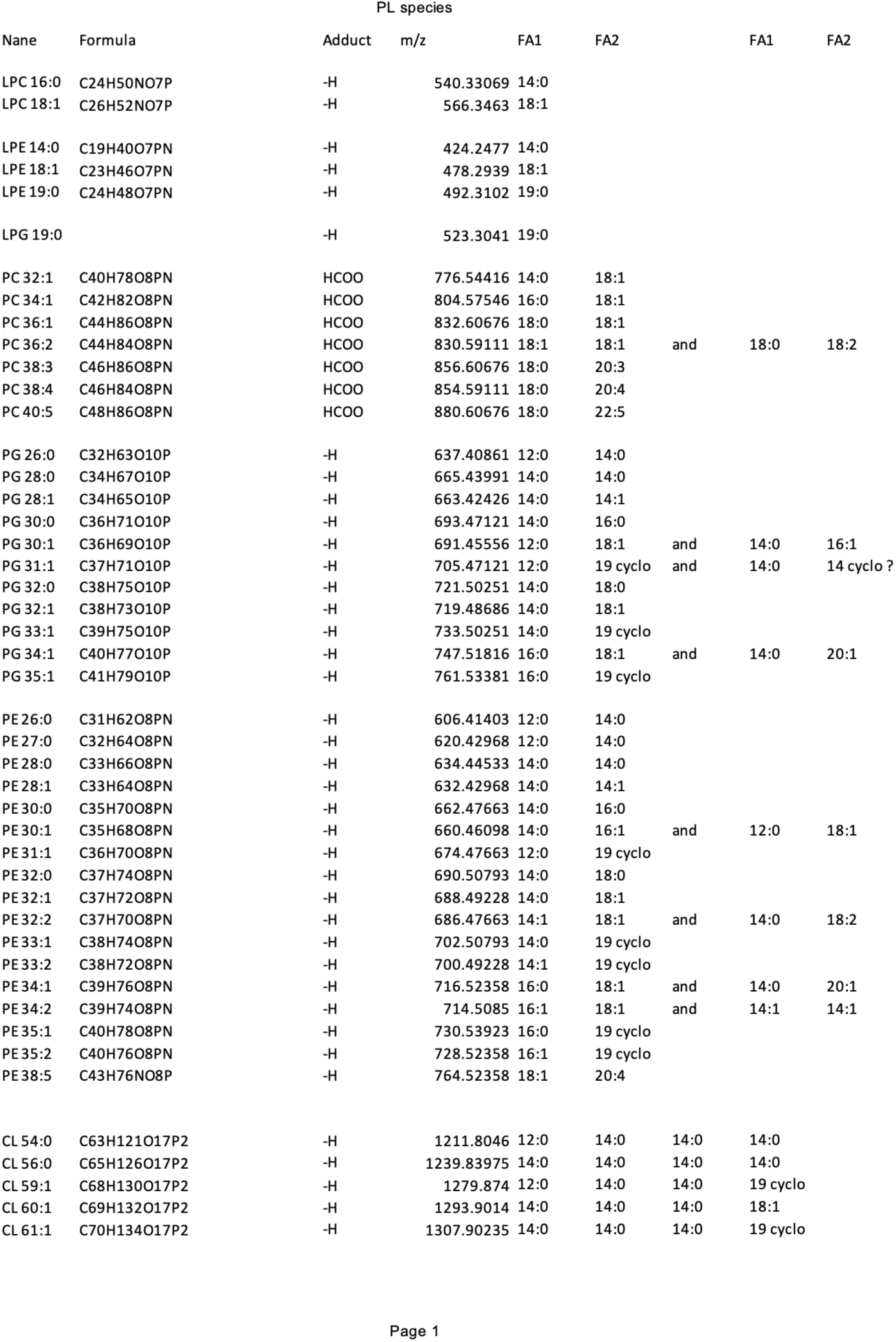
Phospholipid species in membranes of *H. pylori*.

## References

1. Schulz C, Liou J-M, Alboraie M, Bornschein J, Campos Nunez C, Coelho LG, et al. *Helicobacter pylori* antibiotic resistance: a global challenge in search of solutions. Gut. 2025;74: 1561–1570. doi:10.1136/gutjnl-2025-335523

2. Sati H, Carrara E, Savoldi A, Hansen P, Garlasco J, Campagnaro E, et al. The WHO Bacterial Priority Pathogens List 2024: a prioritisation study to guide research, development, and public health strategies against antimicrobial resistance. Lancet Infect Dis. 2025;25: 1033–1043. doi:10.1016/S1473-3099(25)00118-5

3. Rhee K-H, Park J-S, Cho M-J. Helicobacter pylori: bacterial strategy for incipient stage and persistent colonization in human gastric niches. Yonsei Med J. 2014;55: 1453– 1466. doi:10.3349/ymj.2014.55.6.1453

4. Yoshiyama H, Nakazawa T. Unique mechanism of Helicobacter pylori for colonizing the gastric mucus. Microbes and Infection. 2000;2: 55–60. doi:10.1016/S1286-4579(00)00285-9

5. Bonsor DA, Sundberg EJ. Roles of Adhesion to Epithelial Cells in Gastric Colonization by Helicobacter pylori. Adv Exp Med Biol. 2019;1149: 57–75. doi:10.1007/5584_2019_359

6. Nilsson H-O, Blom J, Abu-Al-Soud W, Ljungh A A, Andersen LP, Wadström T. Effect of cold starvation, acid stress, and nutrients on metabolic activity of Helicobacter pylori. Appl Environ Microbiol. 2002;68: 11–19. doi:10.1128/AEM.68.1.11-19.2002

7. Zeng H, Guo G, Mao XH, Tong WD, Zou QM. Proteomic insights into Helicobacter pylori coccoid forms under oxidative stress. Curr Microbiol. 2008;57: 281–286. doi:10.1007/s00284-008-9190-0

8. Cheng T, Mai P-Y, Adedeji-Olulana AF, Smith OER, Wheeler R, Berry J, et al. Morphological transformation in Helicobacter pylori is a dynamic process leading to two types of coccoid. bioRxiv; 2026. p. 2026.01.13.698397. doi:10.64898/2026.01.13.698397

9. Bode G, Mauch F, Malfertheiner P. The coccoid forms of Helicobacter pylori. Criteria for their viability. Epidemiol Infect. 1993;111: 483–490. doi:10.1017/s0950268800057216

10. Loke MF, Ng CG, Vilashni Y, Lim J, Ho B. Understanding the dimorphic lifestyles of human gastric pathogen Helicobacter pylori using the SWATH-based proteomics approach. Sci Rep. 2016;6: 26784. doi:10.1038/srep26784

11. Sörberg M, Nilsson M, Hanberger H, Nilsson LE. Morphologic conversion of Helicobacter pylori from bacillary to coccoid form. Eur J Clin Microbiol Infect Dis. 1996;15: 216–219. doi:10.1007/BF01591357

12. Cheng T, Boneca IG. The shapeshifting Helicobacter pylori: From a corkscrew to a ball. Mol Microbiol. 2024;121: 260–274. doi:10.1111/mmi.15218

13. Buddelmeijer N. The molecular mechanism of bacterial lipoprotein modification— How, when and why? FEMS Microbiology Reviews. 2015;39: 246–261. doi:10.1093/femsre/fuu006

14. McClain MS, Voss BJ, Cover TL. Lipoprotein Processing and Sorting in Helicobacter pylori. mBio. 2020;11: e00911–20. doi:10.1128/mBio.00911-20

15. Jung MS, Piazuelo MB, Brackman LC, McClain MS, Algood HMS. Essential role of Helicobacter pylori apolipoprotein N-acyltransferase (Lnt) in stomach colonization. Infect Immun. 2023;91: e0036923. doi:10.1128/iai.00369-23

16. Legood S, Oliveira Paiva A, Taib N, Ruffiot T, Gribaldo S, Gomperts Boneca I, et al. Arm and head domain in highly conserved lipoprotein modification enzyme Lgt determine functional diversity among bacterial pathogens. mBio. 2026;0: e00600–26. doi:10.1128/mbio.00600-26

17. Mao G, Zhao Y, Kang X, Li Z, Zhang Y, Wang X, et al. Crystal structure of E. coli lipoprotein diacylglyceryl transferase. Nat Commun. 2016;7: 10198. doi:10.1038/ncomms10198

18. Singh W, Bilal M, McClory J, Dourado D, Quinn D, Moody TS, et al. Mechanism of Phosphatidylglycerol Activation Catalyzed by Prolipoprotein Diacylglyceryl Transferase. J Phys Chem B. 2019;123: 7092–7102. doi:10.1021/acs.jpcb.9b04227

19. Pailler J, Aucher W, Pires M, Buddelmeijer N. Phosphatidylglycerol::Prolipoprotein Diacylglyceryl Transferase (Lgt) of Escherichia coli Has Seven Transmembrane Segments, and Its Essential Residues Are Embedded in the Membrane. J Bacteriol. 2012;194: 2142–2151. doi:10.1128/JB.06641-11

20. Rachwalski K, Madden SJ, Ritchie N, French S, Bhando T, Girgis-Gabardo A, et al. A screen for cell envelope stress uncovers an inhibitor of prolipoprotein diacylglyceryl transferase, Lgt, in Escherichia coli. iScience. 2024;27: 110894. doi:10.1016/j.isci.2024.110894

21. Diao J, Komura R, Sano T, Pantua H, Storek KM, Inaba H, et al. Inhibition of Escherichia coli Lipoprotein Diacylglyceryl Transferase Is Insensitive to Resistance Caused by Deletion of Braun’s Lipoprotein. Comstock LE, editor. J Bacteriol. 2021;203. doi:10.1128/JB.00149-21

22. McClain MS, Bryant KN, McDonald WH, Algood HMS, Cover TL. Identification of an Essential LolD-Like Protein in Helicobacter pylori. J Bacteriol. 2023;205: e0005223. doi:10.1128/jb.00052-23

23. Kovacs-Simon A, Titball RW, Michell SL. Lipoproteins of Bacterial Pathogens. Andrews-Polymenis HL, editor. Infect Immun. 2011;79: 548–561. doi:10.1128/IAI.00682-10

24. Senkovich OA, Yin J, Ekshyyan V, Conant C, Traylor J, Adegboyega P, et al. Helicobacter pylori AlpA and AlpB bind host laminin and influence gastric inflammation in gerbils. Infect Immun. 2011;79: 3106–3116. doi:10.1128/IAI.01275-10

25. Ding H, Zeng H, Huang L, Dong Y, Duan Y, Mao X, et al. Helicobacter pylori chaperone-like protein CagT plays an essential role in the translocation of CagA into host cells. J Microbiol Biotechnol. 2012;22: 1343–1349. doi:10.4014/jmb.1202.02025

26. O’Rourke EJ, Pinto AV, Petroni EA, Tolmasky ME, Ielpi L. Evidence for the active role of a novel nuclease from Helicobacter pylori in the horizontal transfer of genetic information. J Bacteriol. 2004;186: 2586–2593. doi:10.1128/JB.186.9.2586-2593.2004

27. Vallese F, Mishra NM, Pagliari M, Berto P, Codolo G, de Bernard M, et al. *Helicobacter pylori* antigenic Lpp20 is a structural homologue of Tipα and promotes epithelial-mesenchymal transition. Biochimica et Biophysica Acta (BBA) -General Subjects. 2017;1861: 3263–3271. doi:10.1016/j.bbagen.2017.09.017

28. Rosinke K, Starai VJ, Hoover TR. Helicobacter pylori HP0018 Has a Potential Role in the Maintenance of the Cell Envelope. Cells. 2024;13: 1438. doi:10.3390/cells13171438

29. Nguyen D, Ivester RG, Rosinke K, Hoover TR. Helicobacter pylori HP0135 Is a Small Lipoprotein That Has a Role in Outer Membrane Stability. Molecules. 2025;30: 204. doi:10.3390/molecules30020204

30. Rosinke K, Tachiyama S, Mrásek J, Liu J, Hoover TR. A Helicobacter pylori flagellar motor accessory is needed to maintain the barrier function of the outer membrane during flagellar rotation. PLoS Pathog. 2025;21: e1012860. doi:10.1371/journal.ppat.1012860

31. Legood S, Seng D, Boneca IG, Buddelmeijer N. A Defect in Lipoprotein Modification by Lgt Leads to Abnormal Morphology and Cell Death in Escherichia coli That Is Independent of Major Lipoprotein Lpp. Maupin-Furlow JA, editor. J Bacteriol. 2022;204: e00164–22. doi:10.1128/jb.00164-22

32. Tsang J, Hoover TR. Basal Body Structures Differentially Affect Transcription of RpoN- and FliA-Dependent Flagellar Genes in Helicobacter pylori. Journal of Bacteriology. 2015;197: 1921–1930. doi:10.1128/jb.02533-14

33. Baca-Porcel Á, Légeret B, Six A, Cabanel M, Le Cossec M, Guegano A, et al. The algal homolog of the plant CER1 and CER3 proteins is a bifunctional hydrocarbon-forming enzyme. New Phytol. 2026;251: 1311–1328. doi:10.1111/nph.71257

34. Suzuki H, Cuiné S, Légeret B, Wijffels RH, Hulatt CJ, Li-Beisson Y, et al. Phosphorus starvation induces the synthesis of novel lipid class diacylglyceryl glucuronide and diacylglyceryl-N,N,N-trimethylhomoserine in two species of cold-adapted microalgae Raphidonema (Chlorophyta). The Plant Journal. 2025;121: e17227. doi:10.1111/tpj.17227

35. McClain MS, Boeglin WE, Algood HMS, Brash AR. Fatty acids of Helicobacter pylori lipoproteins CagT and Lpp20. Microbiol Spectr. 2024;12: e0047024. doi:10.1128/spectrum.00470-24

36. Chu JK, Zhu S, Herrera CM, Henderson JC, Liu J, Trent MS, et al. Loss of a Cardiolipin Synthase in Helicobacter pylori G27 Blocks Flagellum Assembly. J Bacteriol. 2019;201: e00372–19. doi:10.1128/JB.00372-19

37. McClain MS, Shaffer CL, Israel DA, Peek RM, Cover TL. Genome sequence analysis of Helicobacter pylori strains associated with gastric ulceration and gastric cancer. BMC Genomics. 2009;10: 3. doi:10.1186/1471-2164-10-3

38. Colland F, Rain JC, Gounon P, Labigne A, Legrain P, De Reuse H. Identification of the Helicobacter pylori anti-sigma28 factor. Mol Microbiol. 2001;41: 477–487. doi:10.1046/j.1365-2958.2001.02537.x

39. Anselmet M, Xénard L, Albert M, Arias-Cartin R, Hicham S, Pokorny L, et al. Automated Optimization of Bacterial Tracking Pipelines With TrackMate 8. Curr Protoc. 2026;6: e70369. doi:10.1002/cpz1.70369

40. Salama NR, Hartung ML, Müller A. Life in the human stomach: persistence strategies of the bacterial pathogen Helicobacter pylori. Nat Rev Microbiol. 2013;11: 385–399. doi:10.1038/nrmicro3016

41. Wang G, Olczak A, Forsberg LS, Maier RJ. Oxidative stress-induced peptidoglycan deacetylase in Helicobacter pylori. J Biol Chem. 2009;284: 6790–6800. doi:10.1074/jbc.M808071200

42. Hazell SL, Evans DJ, Graham DY. Helicobacter pylori catalase. J Gen Microbiol. 1991;137: 57–61. doi:10.1099/00221287-137-1-57

43. Wells DH, Gaynor EC. Helicobacter pylori initiates the stringent response upon nutrient and pH downshift. J Bacteriol. 2006;188: 3726–3729. doi:10.1128/JB.188.10.3726-3729.2006

44. Seib KL, Srikhanta YN, Atack JM, Jennings MP. Epigenetic Regulation of Virulence and Immunoevasion by Phase-Variable Restriction-Modification Systems in Bacterial Pathogens. Annu Rev Microbiol. 2020;74: 655–671. doi:10.1146/annurev-micro-090817-062346

45. Srikhanta YN, Gorrell RJ, Power PM, Tsyganov K, Boitano M, Clark TA, et al. Methylomic and phenotypic analysis of the ModH5 phasevarion of Helicobacter pylori. Sci Rep. 2017;7: 16140. doi:10.1038/s41598-017-15721-x

46. Patterson SK, He JY, Xu Y, Greene EM, Poznyak Y, Nye MV, et al. Methylation Dynamics in Helicobacter pylori: Exploring Acidic Stress Effects on Epigenetic Acclimation. Microorganisms. 2026;14: 1501. doi:10.3390/microorganisms14071501

47. Patel L, Ailloud F, Suerbaum S, Josenhans C. Single-base resolution quantitative genome methylation analysis in the model bacterium Helicobacter pylori by enzymatic methyl sequencing (EM-Seq) reveals influence of strain, growth phase, and methyl homeostasis. BMC Biol. 2024;22: 125. doi:10.1186/s12915-024-01921-1

48. Patterson SK, He JY, Xu Y, Greene EM, Poznyak Y, Nye MV, et al. Methylation Dynamics in Helicobacter pylori: Exploring Acidic Stress Effects on Epigenetic Acclimation. Microorganisms. 2026;14: 1501. doi:10.3390/microorganisms14071501

49. Ehrmann M. Structure, function, and transport of lipoproteins in Escherchia coli. The Periplasm. 2007. pp. 67–79.

50. Mileykovskaya E, Dowhan W. Cardiolipin-dependent formation of mitochondrial respiratory supercomplexes. Chem Phys Lipids. 2014;179: 42–48. doi:10.1016/j.chemphyslip.2013.10.012

51. Mileykovskaya E, Dowhan W. Cardiolipin membrane domains in prokaryotes and eukaryotes. Biochim Biophys Acta. 2009;1788: 2084–2091. doi:10.1016/j.bbamem.2009.04.003

52. Herrera CM, Demey LM, Ellison CK, Trent MS. A surface-exposed cardiolipin synthase provides an unexpected paradigm for maintaining the Gram-negative outer membrane. Proc Natl Acad Sci U S A. 2026;123: e2524588123. doi:10.1073/pnas.2524588123

53. Schirm M, Soo EC, Aubry AJ, Austin J, Thibault P, Logan SM. Structural, genetic and functional characterization of the flagellin glycosylation process in Helicobacter pylori. Mol Microbiol. 2003;48: 1579–1592. doi:10.1046/j.1365-2958.2003.03527.x

54. Deatherage DE, Barrick JE. Identification of mutations in laboratory-evolved microbes from next-generation sequencing data using breseq. Methods Mol Biol. 2014;1151: 165–188. doi:10.1007/978-1-4939-0554-6_12

55. Ducret A, Quardokus EM, Brun YV. MicrobeJ, a tool for high throughput bacterial cell detection and quantitative analysis. Nat Microbiol. 2016;1: 16077. doi:10.1038/nmicrobiol.2016.77

56. Ershov D, Phan M-S, Pylvänäinen JW, Rigaud SU, Le Blanc L, Charles-Orszag A, et al. TrackMate 7: integrating state-of-the-art segmentation algorithms into tracking pipelines. Nat Methods. 2022;19: 829–832. doi:10.1038/s41592-022-01507-1

57. Dowhan W, Bogdanov M. Eugene P. Kennedy’s Legacy: Defining Bacterial Phospholipid Pathways and Function. Front Mol Biosci. 2021;8: 666203. doi:10.3389/fmolb.2021.666203

58. Sohlenkamp C, López-Lara IM, Geiger O. Biosynthesis of phosphatidylcholine in bacteria. Prog Lipid Res. 2003;42: 115–162. doi:10.1016/s0163-7827(02)00050-4

59. Ferrero RL, Cussac V, Courcoux P, Labigne A. Construction of isogenic urease-negative mutants of Helicobacter pylori by allelic exchange. J Bacteriol. 1992;174: 4212–4217. doi:10.1128/jb.174.13.4212-4217.1992

